# An extension of the *Cx(Co)*^*m*^ model of crossover patterning to account for experimental mortality in *Drosophila melanogaster*

**DOI:** 10.64898/2026.01.07.697413

**Authors:** Spencer Koury, Melika Ghasemi Shiran, Eva Hammonds, Cassidy Schneider, Laurie Stevison

**Author notes:** Co-corresponding authors: Department of Biological Sciences, Auburn University, Room 101 Rouse Life Sciences Building, 120 W Samford Ave, Auburn, AL 36849.

## Abstract

Classic recombination experiments designed to test genetic and environmental treatments do not directly measure crossing-over, instead rates and distribution of meiotic events in F_1_ meiocytes is inferred from genetic markers in F_2_ adults. In *Drosophila melanogaster* females this procedure introduces a substantial “missing data problem” because 75% of meiotic chromatids segregate to polar body nuclei and another 11% are transmitted to inviable F_2_ zygotes which cannot be scored for recombination. To address these sources of uncertainty and bias we extend the *Cx(Co)*^*m*^ model of the data-generating process by assuming: 1) programmed double strand breaks occur as a Poisson point process, 2) crossover maturation is a stationary renewal process, 3) chromosome segregation is random one-half thinning of this process, 4) fertilization by X-versus Y-bearing sperm is mendelian, and 5) egg-to-adult survival is binomially distributed with a rate parameter determined by F_2_ marker alleles. To quantify experimental mortality, we performed egg counts in a 6-point X chromosome testcross and marker-free controls on identical genetic backgrounds under standard laboratory conditions. The 19,927 fly dataset reveals 44% F_2_ experimental mortality, and likelihood ratio tests support a model where 36 of the 44% is due to sex-specific, marker-associated viability defects. Variability in X chromosome genetic lengths with experimental mortality can be simulated and we provide case-control 80% power curves to guide experimental design. We propose that differential mortality should be the *de facto* null hypothesis when comparing F_2_ recombinant fractions and provide probabilistic models of the data-generating process to improve characterization of patterns in F_1_ meiotic events.

## Introduction

Meiosis is the specialized cell division that produces haploid gametes from diploid germline cells in sexually reproducing eukaryotes. Meiotic recombination, exchange of genetic information between homologous chromosomes, occurs after the diploid genome is replicated but before the meiocyte undergoes two rounds of cell division to generate haploid meiotic products (Anderson 1925). Reciprocal exchange between homologs, termed crossing-over, promotes accurate segregation of chromosomes and, as a byproduct, generates new combinations of alleles in meiotic products (Bridges 1916). Therefore, rates and distribution of crossing-over along chromosomes have critical roles in both preventing aneuploidy and shaping genomewide polymorphism in the presence of deleterious mutations (Charlesworth *et al*. 1993; Koehler *et al*. 1996).

There are three general approaches to estimating crossover rates (reviewed in Peñalba and Wolf 2020): 1) cytological analysis of meiocytes, 2) from offspring genotypes in controlled matings or pedigrees, and 3) with linkage disequilibrium in random samples from natural populations. The first method is direct observation of physical manifestation of crossing-over and the third method involves model-based inference with extensive population genetic modeling. However, the second approach is somewhat ambiguous; some experimentalists treat observed offspring frequencies as direct recombination rate estimates, whereas mathematicians have developed probabilistic models to address incompleteness of the same data (cf. Haldane 1919; Crow 1990; Zhao and Speed 1996).

The fundamental challenge in designing recombination testcrosses is not F_2_ sample size or sequencing coverage; but rather that meiotic events of interest occur in developing germaria of F_1_ females and, therefore, are not directly observed (figure 1a,b). Furthermore, because each crossover event involves only two homologs in the four-strand bundle of prophase I, and only one strand will migrate to the oocyte pronucleus at telophase II (with the other three strands included in polar body nuclei), there is only a one-half chance of transmitting evidence of any given crossover to F_2_ progeny (Mather 1935). For example, figure 1b illustrates a 3-strand double exchange which has a 0.25 probability of transmitting the no-exchange strand, a 0.50 probability of transmitting a single exchange strand, and only a 0.25 probability of transmitting the strand carrying evidence of both crossovers.

**Figure 1.**
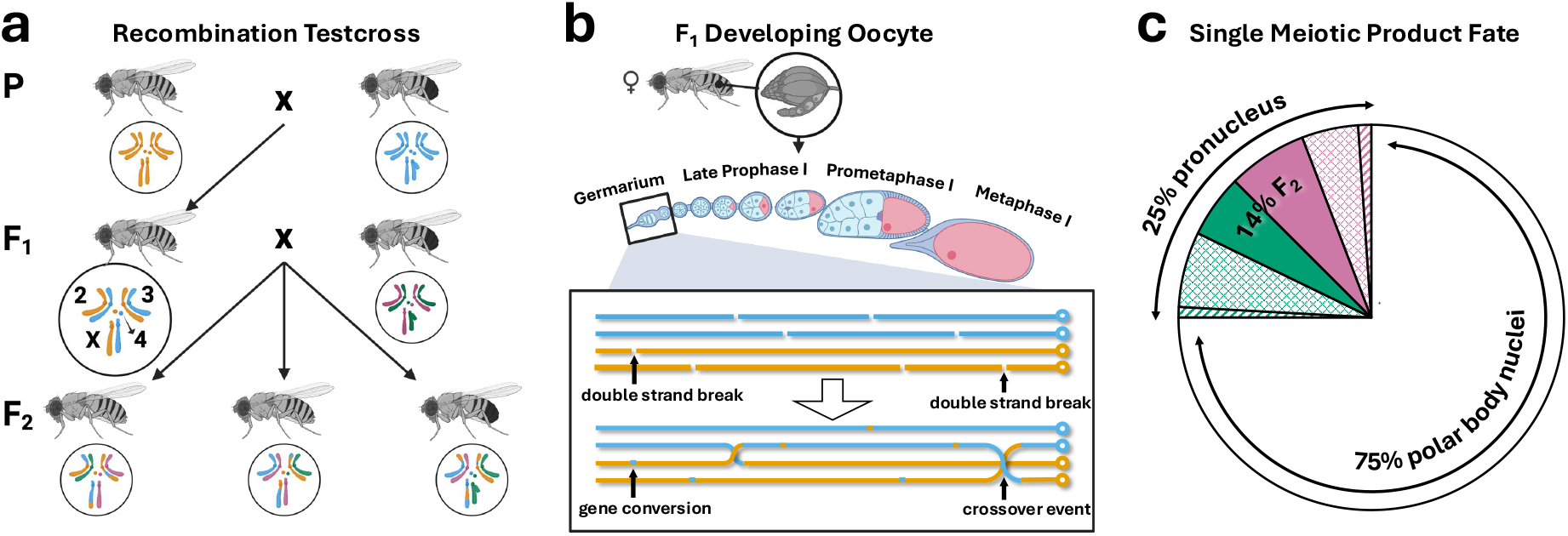
a) *D.melanogaster* recombination testcross illustrated with color-coded karyotypes identifying strain-of-origin for acrocentric X, J-shaped Y, two metacentric autosomes, and the heterochromatic dot chromosomes. The meiosis of interest occurs in heterozygous F_1_ female ovaries but recombination data are collected from F_2_ adult phenotypic classes. b) The F_1_ germarium in context of a single developing ovariole (with inset of enlarged, color-coded, linearized cartoon of the X chromosome prophase I four-strand bundle with 8 double strand breaks repaired as 5 gene conversions and a 3-strand double exchange for reference). Note that programmed double strand breaks, gene conversions, and crossovers occur during prophase I in the germarium long before chromosome segregation that follows metaphase I. c) The fate of single chromatids from the four-strand bundle. 75% of chromatids will segregate to polar body nuclei. Of the 25% of chromatids fated to oocyte pronucleus, one-half are fertilized by X-bearing sperm (pink) and the other half are fertilized by Y-bearing sperm (green). Of these zygotes only 56% will survive to adulthood, with 8% dying for unattributable reason (striped wedges) and 36% dying due to marker-associated viability effects (checked wedges). Thus, overall, there is only a 0.14 probability that any given chromatid from a four-strand bundle will be scored in F_2_ adults for evidence of crossing-over in a recombination testcross (solid-colored pink and green fractions). Created in https://BioRender.com.

Incomplete transmission to the F_2_ generation, however, is not the only difficulty in recombination testcrosses. Our experiments show, on average, only a 56% chance of F_2_ zygotes surviving to adulthood to be scored for recombination (egg-to-adult viability varies based on fertilization by X-versus Y-bearing sperm and markers present in meiotic products). So, given a 3-strand double exchange in the F_1_ germarium (figure 1b), the double crossover detection probability scoring visible markers is 0.56 × 0.25 = 0.14 (figure 1c). While genome sequencing advances might seem to provide solutions (e.g. Miller *et al*. 2016), sequences of meiotic products cannot address this issue, as even infinite sequencing coverage of infinite F_2_ offspring in a marker-free cross has an upper limit of 0.92 × 0.25 = 0.23 probability of detecting such a double crossover event (see Veller *et al*. 2022, for additional limitations of sequencing methods).

To address these fundamental challenges in recombination testcrosses, we extend existing models of the data-generating process as a stochastic function of the underlying meiotic events in F_1_ females and then account for F_2_ experimental mortality with sex-specific viability effects. Briefly, the underlying process-based model states that programmed double strand breaks occur as a homogeneous Poisson point process along the real half-line, a line segment of which defines a chromosome-level genetic map (Haldane 1919). A proportion of such breaks repair as crossover events in a stationary renewal process, with the spacing of these repair events reflecting the strength of crossover interference (Sturtevant 1913; Fisher 1947; Zhao and Speed 1996). Random one-half thinning of this process accounts for asymmetric female meiosis and the one-half chance of sampling evidence of any given reciprocal exchange event (Mather 1935). Fertilization of oocytes by X-versus Y-bearing sperm is mendelian and resulting zygote survival is modeled as a binomial with its rate parameter determined by F_2_ phenotypic class.

The core assumptions in this process-based model were purely theoretical when established by Fisher (1947) and Owen (1949, 1950). They were later re-introduced as Foss *et al*. (1993) “counting model” on biological grounds and then further codified as the “gamma model” (McPeek and Speed 1995), “Poisson-skip model”(Lange *et al*. 1997), and the “chi-square model” (Zhao *et al*. 1995, which is the formulation we extend here). Historically, analysis of recombination testcrosses was solely aimed at building or refining genetic maps (e.g. Haldane 1919; Fisher 1947; Owen 1949; Stam 1979; Crow 1990; Speed 2005). More recently, the underlying models have been applied as mechanistic hypotheses for subcellular meiotic phenotypes to enable statistical inferences about effects of genetic mutations, karyotypic variation, heterochiasmy, and biological aging (e.g. Navarro *et al*. 1997; Zwick *et al*. 1999; Stahl *et al*. 2004; Petkov *et al*. 2007; Zhang *et al*. 2014, 2021; Koury 2023; Kapperud 2025).

Here, we demonstrate that both asymmetric female meiosis and experimental mortality introduce substantial uncertainty and bias when recombination inferences are based only on surviving F_2_ adults. To account for these biases, we extend the *Cx(Co)*^*m*^ model of Zhao *et al*. (1995) to explicitly model zygosis and include sex-specific marker viability effects. The newly presented data is used to generate power curves for case-control experimental designs that can account for stochasticity in crossover patterning, chromosome segregation, fertilization, and experimental mortality. Furthermore, the Appendix outlines how the extended *Cx(Co)*^*m*^ counting model can simulate the sampling distribution for total X chromosome genetic length in *Drosophila melanogaster* and, thereby, allow rigorous null hypothesis statistical testing when comparing genetic or environmental treatments that may alter meiosis but also introduce segregation defects and experimental mortality.

## Results

To investigate sources of experimental mortality in recombination studies, we present a new testcross dataset that includes absolute measures of F_2_ egg-to-adult viability for the model genetic system *D. melanogaster*. We performed two testcrosses using either multiply-marked X chromosomes or a marker-free X chromosome control on identical genetic backgrounds under normal laboratory conditions. Ten replicates of each cross, mated at a density of five females and five males, were serially transferred to generate ten 24-hour broods (figure 2). Immediately following tap transfer, eggs laid were recorded by digital photograph. Thus, when 14,190 F_2_ eclosing adults were scored for visible marker recombination, per vial egg-to-adult survival rates could also be estimated from 19,927 previously imaged F_2_ eggs.

**Figure 2.**
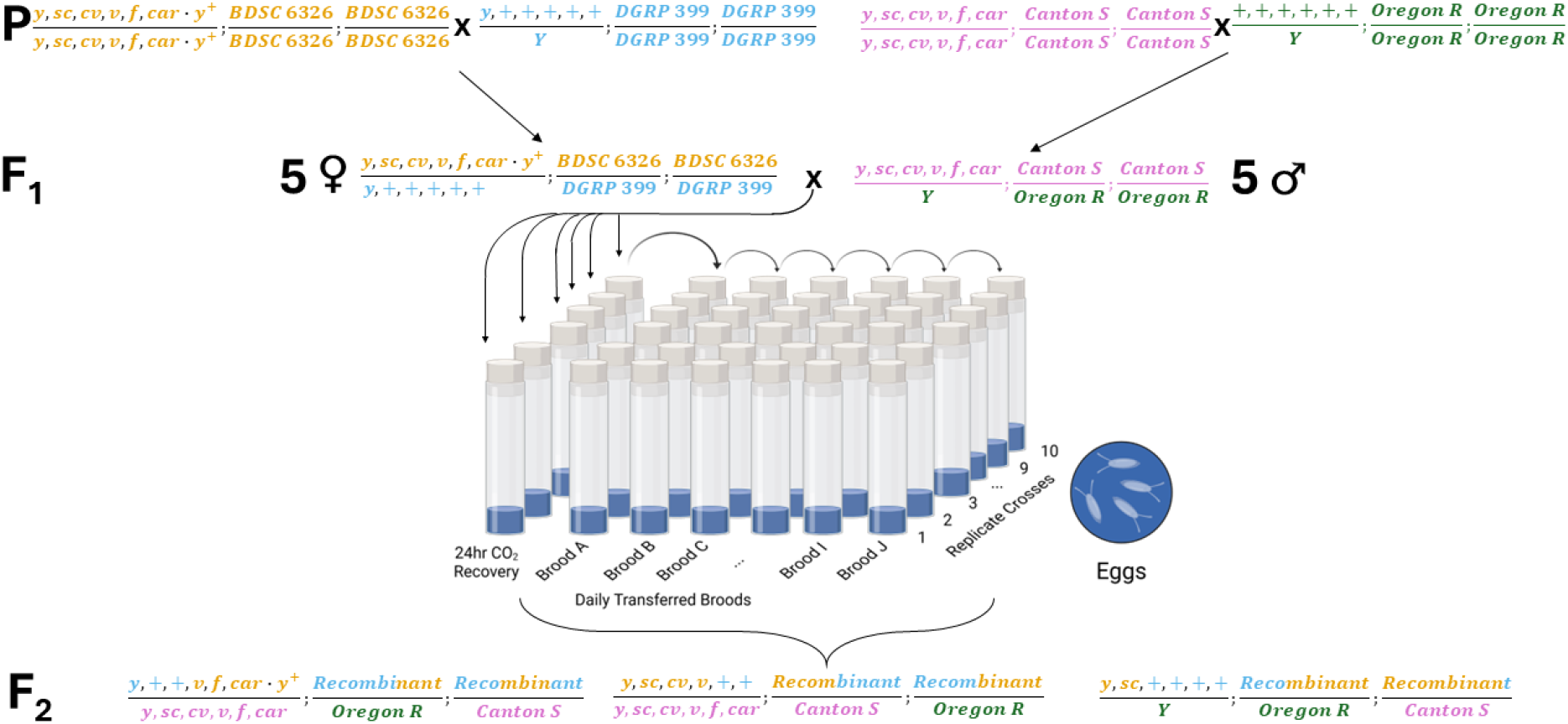
Crossing scheme and experimental design for *D. melanogaster* multiply-marked X chromosome testcross (identical crossing scheme for marker-free controls not shown). Data structure follows the crossing design of ten replicate crosses serially transferred to create ten 24 hour broods. The use of four unique color-coded isogenic strains (yellow *BDSC 6326*, blue *DGRP 399*, pink *Canton-S*, and green *Oregon-R*) in P crosses prevents inbreeding effects from contributing to F_2_ experimental mortality. Detailed marker stock construction can be found in the supplement. Therefore, in a common garden design with the same environmental conditions and identical genetic backgrounds, all excess mortality in multiply-marked crosses versus marker-free controls can be attributed to the presence of *y*^*1*^, *sc*^*1*^, *cv*^*1*^, *v*^*1*^, *f*^*1*^, *car*^*1*^, and *Dp(1;1)sc*^*V1*^, *y*^*+*^. Note *y*^*1*^ is homozygous at its native locus and is not informative, but is required for the use of *Dp(1;1)sc*^*V1*^, *y*^*+*^ as a centromeric marker.

In marker-free crosses, 7.9% of F_2_ eggs failed to develop into F_2_ adults. Assuming a 1:1 primary sex ratio, these failures were evenly distributed among males and females (7.0% vs. 8.7%, *F*^1,188^ = 0.56, *p* = 0.46) and showed no maternal age effects across 10 days (*b*_Y·X_ = −0.006, *t*_s_(188) = −^*1*^ 1.55, *p* = 0.12) (table S1). When X-linked visible markers *y, sc*^*1*^, *cv*^*1*^, *v*^*1*^, *f*^*1*^, *car*^*1*^, and *Dp(1;1)sc*^*V1*^, *y*^*+*^ were segregating on the same genetic background, 44.3% of F_2_ eggs failed to develop into F_2_ adults and could not be scored for recombination. In contrast to marker-free crosses, multiply-marked cross failures were more prevalent in males than females (50.6% vs. 37.9%, *F*_1,128_ = 71.51, *p* = 5.35 × 10^-14^), again with no maternal age effect (*b*_Y·X_ = 0.003, *t*_s_(128) = 1.24, *p* = 0.22) (table S2). Statistically significant replicate cross effects in the presence of markers, but not in marker-free crosses, suggests that markers both increase experimental mortality and also that mortality becomes more variable (table S1 and S2). Without a model of the underlying meiotic processes and explicit treatment of sex-specific marker viability, the degree of experimental mortality in this dataset is consistent with *D. melanogaster* X chromosome genetic map lengths anywhere between 50 and 100 centiMorgans, and thus a model-based approach is necessary to quantify uncertainty and bias in genetic map length estimates.

The F_2_ mortality observed in multiply-marked crosses can be statistically associated with visible markers used to score recombination. To detect uneven recovery of markers in both sexes, we performed single-locus goodness-of-fit G-tests assessing which model best describes viability defects in this dataset. In order of increasing complexity these single-locus mendelian segregation models are:

*H*_*0*_: no excess experimental mortality in multiply-marked crosses above and beyond the marker-free cross,

*H*_*1*_: excess experimental mortality in multiply-marked crosses is random with respect to marker alleles,

*H*_*2*_: excess experimental mortality in multiply-marked crosses is solely due to mutant alleles at marker loci, and

*H*_*3*_: excess experimental mortality in multiply-marked crosses is due to viability effects associated with both mutant and wildtype alleles at marker loci.

The added viability effects in each hypothesis were estimated by the least squares solution to the five-equation systems given in supplemental material. For all markers, the single-locus goodness-of-fit to *H*^*0*^, *H*^*1*^, and *H*^*2*^ were rejected (table 1, tables S3-S8). Thus, full viability models (*H*_*3*_ estimating four parameters from the data with zero degrees of freedom) are necessary to describe experimental mortality in even recombination testcrosses with no experimental treatments. For all six markers, the best fit model estimated viability effects of both mutant and wildtype alleles that differ significantly between the sexes (table S3, figure S2).

**Table 1.** Above the line are single locus goodness-of-fit for *H*_*0*_, *H*_*1*_, *H*_*2*_ reporting the G statistic and denoting statistical with *** to indicate *p* < 0.001 for all G-tests. Below the line are the zero degree of freedom viability estimates from the full viability model *H*_*3*_.

| Model | df<sup>a</sup> | sc<sup>b</sup> | cv<sup>c</sup> | v<sup>d</sup> | f<sup>e</sup> | car<sup>f</sup> | Dp(1;1)sc<sup>VI</sup> g |
| --- | --- | --- | --- | --- | --- | --- | --- |
| $H_0$ | 4 | 10599*** | 10965*** | 10646*** | 10501*** | 10487*** | 10484*** |
| $H_1$ | 3 | 256*** | 622*** | 303*** | 159*** | 144*** | 141*** |
| $H_2$ | 2 | 1063*** | 641*** | 988*** | 1289*** | 1331*** | 1342*** |
| $H_3$ | $v_{\text{female wildtype}}$ | 0.77 | 0.82 | 0.77 | 0.74 | 0.74 | 0.74 |
| | $v_{\text{male wildtype}}$ | 0.66 | 0.75 | 0.67 | 0.62 | 0.61 | 0.61 |
| | $v_{\text{female mutant}}$ | 0.59 | 0.54 | 0.58 | 0.62 | 0.62 | 0.62 |
| | $v_{\text{male mutant}}$ | 0.43 | 0.34 | 0.41 | 0.47 | 0.48 | 0.48 |

<sup>a</sup> see supplemental material for definition of viability parameters estimated from the data<sup>b</sup> scute, X:396,060..397,497 all genomic locations as per Release 6 plus ISO1 MT, FB\_2025\_04<sup>c</sup> crossveinless, X:5,690,002..5,693,084<sup>d</sup> vermilion, X:10,923,972..10,925,631<sup>e</sup> forked, X:17,232,942..17,280,964<sup>f</sup> carnation, X:19,566,019..19,569,401<sup>g</sup> intrachromosomal duplication of roughly X:0..397,000 onto X chromosome right arm

However, the six marker loci reported in table 1 do not segregate independently. Therefore, single-locus tests confound each other and distort viability estimates by a factor related to the genetic distances we wish to estimate. We address this problem by simultaneously estimating multi-locus marker viability effects, genetic lengths, and crossover interference through maximizing the likelihood of observed F_2_ phenotypic class counts under the extended *Cx(Co)*^*m*^ model. Table 2 reports maximum likelihood estimates of the genetic lengths for our full dataset (i.e. pooling all multiply-marked crosses and broods) under the multilocus viability models corresponding to *H*^*0*^, *H*^*1*^, *H*^*2*^, and *H*^*3*^ (see table S9 and figure S3 for multilocus viability estimates).

**Table 2.** Genetic lengths estimated from multilocus dataset for different deterministic map functions or stochastic process *Cx(Co)*^*m*^.

| Origin of Estimates | $X^a$ | $sc - cv$ | $cv - v$ | $v - f$ | $f - car$ | $car - Dp(1;1)sc^{VI}$ | $\ell^b$ | $\theta_i^c$ | LR | df | p-value |
| --- | --- | --- | --- | --- | --- | --- | --- | --- | --- | --- | --- |
| Comeron et al.'s (2012) Map <sup>d</sup> | 66.3 | 14.4 | 19.8 | 23.3 | 4.6 | 4.2 |  |  |  |  |  |
| Haldane's Map Function <sup>e</sup> | 97.6 | 21.3 | 27.3 | 37.9 | 6.3 | 4.8 |  |  |  |  |  |
| Kosambi's Map Function <sup>f</sup> | 80.7 | 18.1 | 22.4 | 29.6 | 6.0 | 4.6 |  |  |  |  |  |
| extended $Cx(Co)^m H_0$ | 76.5 | 17.3 | 21.0 | 27.5 | 6.1 | 4.6 | 33136 | 6 | | | |
| extended $Cx(Co)^m H_1$ | 76.5 | 17.3 | 21.0 | 27.5 | 6.1 | 4.6 | 27927 | 8 | 10418 | 2 | $< 1.0 \times 10^{-325}g$ |
| extended $Cx(Co)^m H_2$ | 78.8 | 17.6 | 21.9 | 28.3 | 6.3 | 4.7 | 27692 | 18 | 470 | 10 | $5.5 \times 10^{-95}h$ |
| extended $Cx(Co)^m H_3$ | 76.8 | 17.0 | 21.3 | 27.8 | 6.1 | 4.6 | 27633 | 30 | 118 | 12 | $7.1 \times 10^{-20}i$ |

<sup>a</sup> Full X chromosome map consisting of 5 listed intervals
<sup>b</sup> Log-likelihood of the probabilistic model denoted as $\ell$
<sup>c</sup> The number of parameters in the model denoted as $\theta_i$
<sup>d</sup> Comeron map values included for comparison, these data do not come from classic recombination testcrosses
<sup>e</sup> Haldane's map function applied to raw recombinant fractions from recombination testcrosses presented here
<sup>f</sup> Kosambi's map function applied to raw recombinant fractions from recombination testcrosses presented here
<sup>g</sup> Likelihood ratio test of $H_2$ vs. $H_1$
<sup>h</sup> Likelihood ratio test of $H_2$ vs. $H_1$
<sup>i</sup> Likelihood ratio test of $H_3$ vs. $H_2$

Because the multi-locus versions of *H*^*0*^, *H*^*1*^, *H*^*2*^, and *H*^*3*^ can be strictly nested by constraining specified viability effects to unity (see Appendix), likelihood ratio tests can evaluate improved fit relative to increasing complexity of viability models. Likelihood ratio tests reveal the added parameters modeling experimental mortality were well-justified (*H*_1_ vs. *H*_0_: *LR* = 10418, *d f* = 2, *p <* 1.0 × 10^−325^), excess mortality was not random with respect to marker alleles (*H*_2_ vs. *H*_1_: *LR* = 470, *d f* = 10, *p* = 5.5 × 10^−95^), and both mutant and wildtype alleles at marker loci had sex-specific viability effects (*H*_3_ vs. *H*_3_: *LR* = 118, *d f* = 12, *p* = 7.1 × 10^−20^). Similar to the single locus tests, multilocus analysis supports *H*_*3*_ as the best viability model for the data-generating process.

To explore variability in genetic lengths, we fit the extended *Cx(Co)*^*m*^ model to data collected from single vials as the experimental unit. The observed variation was then partitioned between individual crosses and broods as fixed categorical effects with a Type I ANOVA (table S10). There was no significant effect of cross (*F*^6,54^ = 1.10, *p* = 0.34) or brood (*F*^9,54^ = 1.04, *p* = 0.42) on genetic length. Finally, improvement in statistical power can be achieved by pooling the data from multiple crosses or by broods (table S11 and S12). Figure 3 illustrates the 80% power curves in a case-control design with respect to the F_1_ experimental units of analysis under the different data pooling strategies.

**Figure 3.**
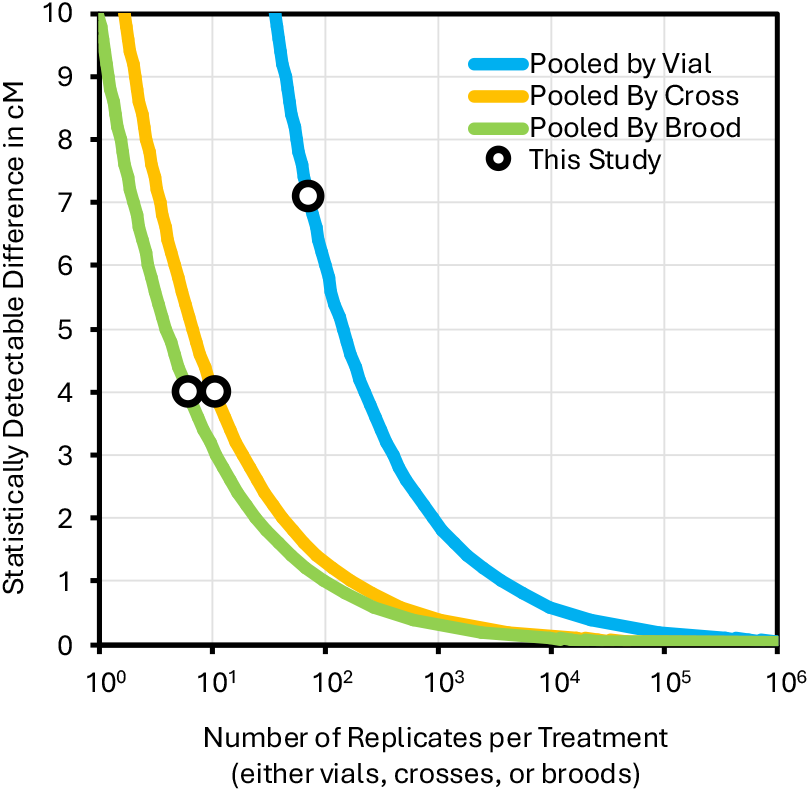
Case-control design 80% power curves for comparison of X chromosome genetic lengths under different pooling schemes with respect to the F_1_ experimental units of analysis (data pooling by either vials, crosses, or broods)

## Discussion

The physical locations of meiotic crossing-over along the chromosome axis are of fundamental importance to a wide range of genetical research from cellular biology to evolutionary theory. Here, we show classic methods for characterizing the rate and distribution of crossing-over has a substantial missing data problem. In female-limited recombination of *D. melanogaster*, 75% of meiotic chromatids (and the accompanying evidence of meiotic events) are lost to polar body nuclei in F_1_ females (figure 1c). Another 11% of chromatids are lost in F_2_ zygotes that die before being scored for recombination (figure 1c). A further 5% of chromatids carry no evidence of recombination even though their four-strand bundle of origin may have had multiple exchanges (figure 1b). Therefore, expressing recombination rates as a fraction:

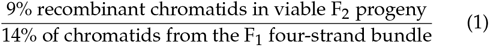

approximates a 66 cM X chromosome but reveals little about the rate and distribution of actual meiotic events in prophase I of F_1_ females.

While the missing data problem outlined above causes considerable uncertainty in estimates, perhaps more concerning is that both numerator and denominator in recombinant fractions are biased by viability effects of the visible markers scored for recombination. We show that experimental mortality (44%) is using viability models *H*_*0*_, *H*_*1*_, *H*_*2*_, and *H*_*3*_. Likelihood ratio tests can only be conducted on probabilistic model-based methods. the major signal in recombination studies, and the majority of this death (36 of the 44%) is due to sex-specific viability effects associated with both mutant and wildtype alleles. Thus, simple calculation of recombination rates from the observed F_2_ adults can be strongly biased (table 1, 2, S3-S9) and prohibits comparison of rates between experimental genotypes and/or treatments. To address this problem, we model the data-generating process by extending the *Cx(Co)*^*m*^ model to a system of 129 equations defining the probability of observing an F_2_ adult fly in any of 64 sex-specific phenotypic classes as well as the number of inviable F_2_ zygotes that could not be scored for recombination (see Appendix). This increased model complexity (24 additional parameters for our experiments) is well-justified by likelihood ratio tests and suggests that even more nuanced analyses of recombination datasets (e.g. McPeek and Speed 1995; Lange *et al*. 1997; Housworth and Stahl 2003) can be substantially improved by incorporating both measures and models of F_2_ survival.

Experimental mortality in *D. melanogaster* genetic studies has long been recognized, but is often overlooked in the design and analysis of recombination testcrosses (but see Bridges and Olbrycht 1926; Redfield 1957; Hayman and Parsons 1962; Koury 2023). The present study builds on a published protocol for quantifying this source of missing data Schneider *et al*. (2025), corrects biases that differential mortality introduces to experimental observations, and can ultimately incorporate appropriate uncertainty to the maximum likelihood estimates through mechanistic simulation of sampling distributions (see Appendix).

The newly presented *D. melanogaster* recombination dataset, and analysis with the extended *Cx(Co)*^*m*^ counting model, demonstrates that F_2_ experimental mortality is overwhelmingly due to sex-specific viability effects associated with both mutant and wildtype alleles even under well-controlled, benign crossing conditions. Future recombination studies, especially those introducing strong genetic or environmental perturbations to meiosis, should directly measure experimental mortality as a likely source of substantial uncertainty and bias. Therefore, we propose the *de facto* null hypothesis in recombination studies be that changes in F_2_ marker distribution results from experimental mortality and that analysis with probabilistic models of the data-generating process are essential for valid statistical inference about rates and distributions of meiotic events in F_1_ females.

## Materials and methods

Both multiply-marked and marker-free X chromosome crosses were conducted in a common garden design (i.e. same environmental conditions and identical genetic backgrounds). Multiply-marked X chromosome consisted of seven visible markers *y*^*1*^, *sc*^*1*^, *cv*^*1*^, *v*^*1*^, *f*^*1*^, *car*^*1*^, and *Dp(1;1)sc*^*V1*^, *y*^*+*^ in a stock reconstituted from Bloomington *Drosophila* Stock Center lines (BDSC 177, 1515, and 4914) (Grell 1978). This reconstituted multiply-marked chromosome was then background-replaced with autosomes of BDSC 6326 using a balancer-translocation stock derived from BDSC 2475 (*w*\*; *T(2;3)ap*^*Xa*^*/In(2L)Cy, In(2R)Cy; TM3, Sb*^*1*^).

Centromeric marker *Dp(1;1)sc*^*V1*^, *y*^*+*^ required introgression of *y*^*1*^ from BDSC 1515 into *Drosophila melanogaster* Genetic Reference Panel (DGRP) line 399 (BDSC 25192) by repeatedly back-crossing for >10 generations. Thus, all F_1_ experimental females in multiply-marked testcrosses were whole autosome heterozygotes for BDSC 6326 and DGRP 399, whole chromosome heterozygotes for the multiply-marked X chromosome with a homozygous *y*^*1*^ at its native locus (figure 2). F_1_ experimental females were crossed to whole genome heterozygotes for Oregon-R (BDSC 25211) and Canton-S (BDSC 64349) carrying visible markers *y*^*1*^, *sc*^*1*^, *cv*^*1*^, *v*^*1*^, *f*^*1*^, and *car*^*1*^ (figure 2). Marker-free crosses involved the same stocks to reproduce identical genetic backgrounds but with no X-linked markers (i.e. pure strains of Canton-S, Oregon-R, DGRP 399, and BDSC 6326 with *w*^*1118*^ allele replaced by *w*^*+*^). See supplemental material for detailed stock construction notes and crossing diagrams.

Crosses were conducted in polystyrene vials with standard cornmeal *Drosophila* media containing 0.01 v/v McCormick blue food coloring to increase egg contrast in digital pictures. Additionally, 50 *µ*L of live yeast paste (0.2 g/mL) with 0.02 v/v blue food coloring was added to induce egg laying behavior (Schneider *et al*. 2025). Vials were housed in a Percival incubator at 21°, 70% relative humidity, and 12:12 hour light-dark cycle.

Virgin F_2_ flies were collected over 72-hours, aged an additional 72 hours, then crossed at densities of five females to five males under light CO_2_ anesthesia. After 24-hour recovery, mated flies were tap transferred onto fresh food establishing “Brood A”. Every subsequent 24-hours for 10 days, flies were tap transferred onto freshly prepared food to generate new broods (Broods B-J). Immediately after tap transferring flies, food surface was imaged for egg counting (Schneider *et al*. 2025). 72 hours after imaging, a 15 cm^2^ piece of Kimwipe was inserted to food and wetted with 100 *µ*L of 0.01M propionic acid/0.001M phosphoric acid.

Starting 10 days after egg laying and continuing until all viable adults had eclosed, sex-specific adult phenotypic classes were scored for recombination. Logistical complications prevented scoring multiply-marked crosses in broods G, H, and I. Two independent digital egg counts were performed; because digital scoring consistently undercounts eggs per vial and individual observers differ in their undercounting, both egg counts were transformed using standard curves constructed from independent dataset of Schneider *et al*. (2025) (figure S1). Number of F_2_ eggs failing to develop into F_2_ adults was calculated as the difference between the average of transformed egg counts and the number of F_2_ adults scored for recombination.

Single-locus viability effects were tested using five F_2_ adult classes (female wildtype, female mutant, male wildtype, male mutant, and lethal zygotes) independently for the six marker loci. The added viability effects in each hypothesis (*H*^*0*^, *H*^*1*^, *H*^*2*^, and *H*_*3*_) was estimated by the least squares solution to respective five-equation systems given in supplemental material. Single-locus goodness-of-fit was assessed with G-tests (Sokal and Rohlf 1995).

Multi-locus viability effects and genetic lengths were estimated by maximizing the likelihood of 129 F_2_ class counts (2^6^ phenotypic classes × 2 sexes + 1 for lethal zygotes) under the extended *Cx(Co)*^*m*^ counting model for the full dataset (pooling all vial counts). Custom R scripts to calculate the likelihood of the data under hypotheses *H*^*0*^, *H*^*1*^, *H*^*2*^, and *H*^*3*^ as well as maximizing likelihoods using the downhill simplex method are available through GitHub (see Data availability section). Likelihood maximization was implemented with R package “dfoptim” where all viability estimates were constrained between 0 and 1 (Varadhan *et al*. 2023).

To characterize the patterns of variability in the estimated genetic lengths, we also fit the extended *Cx(Co)*^*m*^ counting model to data with single vials, pooled broods, and pooled crosses as the experimental units. The observed variation was partitioned between crosses and/or broods as fixed effects with a Type I ANOVA for each pooling scheme (table S10, S11, S12). Case-control 80% power curves were determined by “Number of replications needed to detect a given ‘true’ difference between two means” method of Sokal and Rohlf (1995) using:

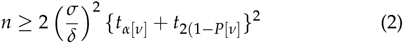

where *n* is the number of replications, *σ* is ‘true’ standard deviation estimated from respective ANOVA tables, *δ* is minimal detectable difference, *ν* is degrees of freedom in the sample standard deviation, *α* is significance level, *P* is intended power of the test, with the values of *tα*[*ν*] and *t*2(1 – *P*[*ν*] coming from two tailed t-tables.

## Data availability

Strains are available upon request. File S1 contains detailed strain construction notes as well as descriptions of all supplemental materials. Raw count data, R code for all analyses, and further documentation of model structure and fitting procedures can be found at https://github.com/StevisonLab/CxCoM-with-Viability.

## Acknowledgments

We would like to thank members of the Stevison Lab and Dr. Vince Ficarrotta for discussion during the development of this project.

