## Appendix for "An extension of the *Cx(Co)*^*m*^ model of crossover patterning to account for experimental mortality in *Drosophila melanogaster*"

### Appendix: Extended *Cx(Co)$^m$* model of crossover patterning

The counting model of crossing-over refers to a family of stochastic processes that share common assumptions in describing the patterning of genetic exchange along meiotic chromosomes (Fisher 1947; Owen 1949, 1950; Foss *et al.* 1993; McPeek and Speed 1995; Zhao *et al.* 1995b; Lange *et al.* 1997; Copenhaver *et al.* 2002). The chi-square $Cx(Co)^m$ formulation of the counting model proposes crossover precursors $(C)$ occur as a homogenous Poisson point process along the genetic map, some crossover precursors mature into crossover events $(Cx)$ in a stationary renewal process, and and the remaining crossover precursors are designated non-crossover events $(Co)$. All three events have straight-forward biological interpretations: crossover precursors $C$ are programmed double strand breaks, crossover $Cx$ events are meiotic repair of double strand breaks resulting in reciprocal genetic exchange between homologs, and non-crossover $Co$ events are meiotic repair of double strand breaks resulting in directional genetic exchange between homologs (i.e. non-crossover gene conversion events). Intuitively, the spacing of crossover $Cx$ events with $m$ intervening non-crossover $Co$ events reflects the strength of crossover interference. Zhao *et al.* (1995b) provide a matrix algebra notation for this model and, where possible, we preserve their naming conventions.

Here, we extend the $Cx(Co)^m$ model to include both random and marker-associated, sex-specific viability effects. This extension requires explicit modeling of fertilization as well as expanding the system of equations to solve for different probabilities of observing reciprocal recombinant classes both within and between the sexes. We introduce the model and our extension by sequentially defining: 1) the probabilities of all possible crossovers in a single interval on the four-strand bundle during prophase I of the $F_1$ developing oocyte, 2) the probabilities of all possible crossover patterns in two adjacent intervals of the four-strand bundle in prophase I, 3) the probabilities of any recombination pattern on a single strand after chromosome segregation in anaphase I and II, 4) the probability of fertilization by X-bearing or Y-bearing sperm, and 5) the survival probability of the resulting zygote as a function of the $F_2$ genotype. This produces a probabilistic model of the data-generating process that can be represented by a system of equations, one for each observable $F_2$ phenotypic class plus one for lethal zygotes. Given a recombination dataset with an absolute measure of experimental mortality, the maximum likelihood solution to this system of equations can be used to estimate parameters in the underlying model(s) of meiosis.

#### Probability of crossover pattern on four-strand bundle

Let $y$ be the rate of crossover precursor $C$ formation along the real half-line. One can think of the chromosome level genetic map as a line segment of the real half-line where this point process takes place with the centromere serving as the origin. Assuming these events are Poisson distributed along the meiotic chromosome means the chance of $s$ number of crossover precursors on the four-strand bundle is:

$$\frac{e^{-y}y^{s}}{s!}. \tag{1}$$

The fraction of precursor $C$ events that mature to crossover $Cx$ events is $1/p$. If programmed double strand breaks occur randomly *among* the four strands in prophase I and there is no chromatid interference (Zhao *et al.* 1995a), then each strand has a one-half chance of being involved in a crossover event (Mather 1935).

However, in the renewal process that governs crossover maturation, the $1/p$ precursor $C$'s resolving as $Cx$ crossover events are not randomly distributed *along* the four-strand bundle. This is because the $Cx(Co)^m$ model assumes a directional and deterministic pattern of a $Cx$ followed by $m$ number of $Co$'s and then a $Cx$ followed by another $m$ number of $Co$'s and so on (thus the nomenclature of $Cx(Co)^m$ counting model). Importantly, this renewal process is also assumed to be stationary, such that the first precursor $C$ event has an equal chance of resolving as a crossover $Cx$ event or any of the $m$ number of sequential non-crossover $Co$ events. This second assumption reintroduces a probabilistic element into the deterministic renewal process. Following Zhao *et al.* (1995b), the occurrence of $k$ crossovers between two markers $l_1$ and $l_2$ (i.e. a single interval) can be computed as:

$$\frac{e^{-y}}{p}\sum_{i=1}^{p}\sum_{j=0}^{p-1}\frac{y^{\,p\,k-p+i+j}}{(p\,k-p+i+j)!}. \tag{2}$$

Equation 2's double summation for the first interval, denoted by subscript one, can conveniently be rewritten in matrix notation as:

$$\frac{1}{p}\,\mathbf{1}\,\boldsymbol{D}_{k_1}(y_1)\,\mathbf{1'}. \tag{3}$$

Where $\mathbf{1}$ is a $1\times p$ vector, $\mathbf{1'}$ is its transpose, and $\boldsymbol{D}_{k_1}(y_i)$ is the $p\times p$ matrix with $i,j^{\text{th}}$ entry $e^{-y_1}y_1^{\,p\,k_1-p+i+j}/(p\,k_1-p+i+j)!$.

Within the first marker-defined interval, the stationarity of the renewal process means a random starting point in the $Cx(Co)^m$ sequence corresponds, in aggregate, to a random stopping point in this sequence irrespective of interval length. Nonetheless, the directionality of this process dictates that the *realized* stopping point of the $Cx(Co)^m$ sequence in the first interval for any given chromosome of a specific oocyte will fully determine the starting point in the adjacent second interval on that chromosome. Zhao *et al.* (1995b) provide a proof that the chance of $k_1$ crossovers between markers $l_1$ and $l_2$ with $k_2$ crossovers in the adjacent interval (between markers $l_2$ and $l_3$) is:

$$e^{-y_2} \sum_{i=1}^{p} p_{k_1}^{p+1-i} \sum_{j=0}^{p-1} \frac{y_2^{p\,k_2-p+i+j}}{(p\,k_2-p+i+j)!}. \tag{4}$$

Rewriting equation 4 in matrix notation, with subscripts 1 and 2 denoting first and second intervals, and the definition of $(p_{k_1}^1 p_{k_1}^2 p_{k_1}^3 ... p_{k_1}^p) = \frac{1}{p}\mathbf{1}\boldsymbol{D}_{\boldsymbol{k_1}}(y_1)$ yields:

$$(p_{k_1}^1\ p_{k_1}^2\ p_{k_1}^3\ ...\ p_{k_1}^p)\ \boldsymbol{D}_{\boldsymbol{k_2}}(y_2)\ \mathbf{1'}. \tag{5}$$

Here, we see that termination of $Cx(Co)^m$ sequence in interval 1 corresponds to the start of the renewal process in interval 2. Making further use of Zhao *et al.*'s (1995b) definition of $(p_{k_1}^1\ p_{k_1}^2\ p_{k_1}^3\ ...p_{k_1}^p)$ allows matrix algebra to account for all possible $Cx(Co)^m$ sequence starting and ending points both *within* an interval and *across* two adjacent intervals of any size as:

$$\frac{1}{p}\ \mathbf{1}\ \boldsymbol{D}_{\boldsymbol{k_1}}(y_1)\ \boldsymbol{D}_{\boldsymbol{k_2}}(y_2)\ \mathbf{1'}. \tag{6}$$

This approach can be extended to any number of adjacent intervals, or even non-adjacent intervals separated by a known genetic distance (e.g. a centromere or structural rearrangement).

#### Probability of recombination pattern on a single strand

Determining the probability of any crossover pattern on the *four-strand* bundle in prophase I is the primary goal of modeling meiotic recombination. Unfortunately, experimental observation of this crossover pattern requires labor-intensive cytological analysis. Therefore, to employ maximum likelihood methods for recombination testcross data we must define the probabilities of any recombination pattern on a *single strand*.

Recall that each crossover event involves only two homologs in the four-strand bundle of prophase I, and only one strand will migrate to the oocyte pronucleus at telophase II (with the other three strands included in polar body nuclei, see figure 1b in main text). Consequently, there is only a one-half chance of transmitting evidence of any given crossover to $F_2$ progeny (Mather 1935). This means the average number of crossover $Cx$ events on each strand given $s$ number of precursor $C$ events is $s/2p$, which defines the total chromosome map length as $100y/2p$ centiMorgans (cM).

However, modeling the specific patterns of crossing-over that produced the $100y/2p$ cM genetic length, rather than the average, requires random one-half thinning of the stochastic point process defined in the first section. Therefore, the observation of no recombination $(N_z)$ in interval $z$ is the result of either:

1) no precursor $C$ events occurring in the interval $z$ $(s = 0)$, denoted $\boldsymbol{D}_{\mathbf{0}}(y_z)$,

2) some precursor $C$ events $(s \geq 1)$, but none that mature into crossover $Cx$ events,

3) some precursor $C$ events $(s \geq 1)$ with some crossover $Cx$ events, but none that involve the single strand migrating to the functional egg pole, or

4) some precursor $C$ events $(s \geq 1)$ maturing to an even number of crossover $Cx$ events that involve the same strand from the four-strand bundle.

In sum, all the ways of observing no recombination in interval $z$ is given by Zhao *et al.*'s (1995b) Appendix Theorem 1:

$$\boldsymbol{N_z} = \boldsymbol{D}_{\mathbf{0}}(y_z)\ + \frac{1}{2} \sum_{s\geq 1} \boldsymbol{D_s}(y_z). \tag{7}$$

In contrast, observation of recombination $(R_z)$ in interval $z$ must necessarily come from $s \geq 1$ precursor $C$ events and either an odd number of crossover $Cx$ events or any even number of crossover $Cx$ events that do not involve the same single strand. Therefore, using Mather's (1935) proof and Zhao *et al.*'s (1995b) Appendix Theorem 1, we relate the observable recombination pattern with the underlying crossover pattern with:

$$\boldsymbol{R_z} = \frac{1}{2} \sum_{s\geq 1} \boldsymbol{D_s}(y_z). \tag{8}$$

At this point our notation diverges from Zhao *et al.* (1995b) who coded recombination data as binary by *interval* and implicitly pooled reciprocal marker classes. Because we are interested in the inequivalence of reciprocal classes due to marker-associated viability effects, our recombination data are coded as binary by *marker*. Thus, a six-point testcross with five intervals formerly had its "probability of recombination interval pattern" coded as $(i_1\ i_2\ i_3\ i_4\ i_5)$ but is here replaced with our "probability of the recombination marker pattern" $(l_1\ l_2\ l_3\ l_4\ l_5\ l_6)$, giving:

$$Pr(l_1\ l_2\ l_3\ l_4\ l_5\ l_6) = \frac{1}{2p}\mathbf{1}\ \boldsymbol{M_1}\ \boldsymbol{M_2}\ \boldsymbol{M_3}\ \boldsymbol{M_4}\ \boldsymbol{M_5}\ \mathbf{1'}, \tag{9}$$

where $\mathbf{M_z} = \mathbf{N_z}$ when $l_z = l_{z+1}$ and $\mathbf{M_z} = \mathbf{R_z}$ when $l_z \neq l_{z+1}$. Please note that the denominator is $2p$ to reflect that our "recombination marker pattern" is the probability of observing only one of two reciprocal classes that result from the same "recombination interval pattern" notation.

Motivated by our experimental dataset from *Drosophila melanogaster* testcrosses, we assume an XY chromosomal sex-determination system where sex chromosome segregation is mendelian. This means there are twice as many $F_2$ phenotypic classes (now including sex denoted $h$) and the probability of the recombination marker pattern $(l_1\ l_2\ l_3\ l_4\ l_5\ l_6\ h)$ for a six-point testcross must be further reduced by a factor of two:

$$Pr(l_1\ l_2\ l_3\ l_4\ l_5\ l_6\ h) = \frac{1}{4p}\mathbf{1}\ \boldsymbol{M_1}\ \boldsymbol{M_2}\ \boldsymbol{M_3}\ \boldsymbol{M_4}\ \boldsymbol{M_5}\ \mathbf{1'}. \tag{10}$$

#### Probability of observing recombination marker patterns

Our notation changes are necessary because sex-specificity of viability effects causes the probability of observing a given recombination marker pattern to no longer be a simple one-half thinning of the crossover patterning process in $F_1$ female meiosis. Sex-specific, marker-associated viability effects make detection probabilities become a function of sex chromosome segregation in $F_1$ *male* meiosis as well as subsequent sex-specific $F_2$ survival rates. To explicitly incorporate both random and marker-associated experimental mortality into the extended $Cx(Co)^m$ model, we reduce the probability of observing a given sex-specific recombination marker pattern by the proportional viability of the marker-free cross as well as each respective marker allele in the multiply-marked cross assuming a multiplicative fitness function.

Comparison of marker-free controls and multiply-marked testcrosses demonstrates that 36 of the 44% experimental mortality is due to sex-specific, marker-associated viability effects. The remaining 8% of zygotic lethality is unattributable to sex, maternal age, or marker effects. The unattributable experimental mortality is: 1) assumed to be random, 2) denoted $1 - v_x$, 3) estimated *a priori* from absolute mortality in marker-free crosses, and 4) treated as a constant during $Cx(Co)^m$ model fitting.

Both goodness-of-fit G-tests and likelihood ratio tests (single-locus and multi-locus datasets, respectively, in the accompanying paper) indicate there were marker-associated viability effects of both mutant and wildtype alleles at all loci, and that these effects differed between the sexes. Therefore, the proportional viability ranging from 0 to 1 are denoted $v_{alh}$ representing the viability effect of allele $a$, at locus $l$, in sex $h$, and the probability of observing recombination pattern $(l_1\ l_2\ l_3\ l_4\ l_5\ l_6)$ in $F_2$ sex $h$ is reduced by:

$$v_{l_1 l_2 l_3 l_4 l_5 l_6 h} = v_x \times v_{a1h} \times v_{a2h} \times v_{a3h} \times v_{a4h} \times v_{a5h} \times v_{a6h}. \quad (11)$$

Using this notation, all four multilocus viability models in the accompanying paper ($H_0$, $H_1$, $H_2$, and $H_3$) can be expressed by constraining one or more $v$ parameters in equation 11 to unity, which also allows straight-forward likelihood ratio testing of these nested hypotheses.

As an example using the full viability model $H_3$, the probability of *forming* a yellow-bodied $F_2$ male zygote with evidence of a single exchange between *vermilion* and *forked* in our study was:

$$Pr(111000m) = \frac{1}{4p}\mathbf{1}\ \boldsymbol{N_1}\ \boldsymbol{N_2}\ \boldsymbol{R_3}\ \boldsymbol{N_4}\ \boldsymbol{N_5}\ \mathbf{1'} = 0.044. \quad (12)$$

However, the probability of *observing* a yellow-bodied $F_2$ male (i.e. surviving egg-to-adulthood development to be scored for recombination) was only:

$$Pr(111000m) = \frac{1}{4p}\mathbf{1}\ \boldsymbol{N_1}\ \boldsymbol{N_2}\ \boldsymbol{R_3}\ \boldsymbol{N_4}\ \boldsymbol{N_5}\ \mathbf{1'}\ v_{111000m} = 0.017. \quad (13)$$

Including the example above, there are 128 unique, sex-specific phenotypic classes in a 6-point testcross. Additionally, the digital record of eggs laid per vial allows calculation of the number of inviable zygotes as the $129^{\text{th}}$ class. Again taking the example of a yellow-bodied $F_2$ male with evidence of a single exchange between *vermilion* and *forked*, the probability of such a fly occurring in our experiment but not surviving to adulthood was:

$$\frac{1}{4p}\mathbf{1}\ \boldsymbol{N_1}\ \boldsymbol{N_2}\ \boldsymbol{R_3}\ \boldsymbol{N_4}\ \boldsymbol{N_5}\ \mathbf{1'}\ (1 - v_{111000m}) = 0.027. \quad (14)$$

Summing across all sex-specific phenotypic classes the total probability of observing an inviable zygote can be expressed as:

$$Pr(inviable) = \sum_a \sum_l \sum_h \frac{1}{4p}\mathbf{1}\boldsymbol{M_1}\boldsymbol{M_2}\boldsymbol{M_3}\boldsymbol{M_4}\boldsymbol{M_5}\mathbf{1'}(1 - v_{l_1 l_2 l_3 l_4 l_5 l_6 h}). \quad (15)$$

While any one phenotypic class's contribution to zygotic lethality is small, together they constitute 44% of the total flies scored in the experiment and must be accounted for in any attempt to infer the rate and distribution of crossing-over in $F_1$ meiosis from the genetic markers of surviving $F_2$ adults.

#### Simulation of sampling distribution

The parameters of the data-generating process and the underlying model of meiosis can be estimated by maximum likelihood. The same data-generating process can be used to simulate the sampling distribution of any summary statistic or meiotic parameter of interest in that dataset. This is particularly useful for estimating confidence intervals as well as statistical comparisons between biological groups. The stated assumptions of the extended $Cx(Co)^m$ model provide a step-by-step guide for this process.

First, the number and spacing of programmed double strand breaks ($C$ events) are simulated along the chromosome as a 1-dimensional Poisson point process on the real half-line representing the genetic map. The rate of this process is given by $Cx(Co)^m$ estimates of $y$. Each $C$ is assigned independently to one of the four strands with equal probability. This procedure explicitly assumes no genetic variation for programmed double strand break placement.

Second, $1/p$ of the $C$ event mature into crossovers ($Cx$ events) following the $Cx(Co)^m$ renewal process using the estimated $m$. This counting process is initiated at the centromere and has a randomly assigned starting point in the $Cx(Co)^m$ sequence to satisfy the stationarity assumption. Template strand for crossover repair is randomly assigned to one of two homologous chromosomes, conditional on the $C$ event strand identity, with equal probability. This procedure explicitly assumes no chromatid interference and no sister chromatid exchange.

Third, one of the four chromatids in the four-strand bundle is randomly selected with equal probability as the meiotic product segregating to the functional egg pole and oocyte pronucleus. This procedure explicitly excludes nondisjunction and assumes no meiotic drive.

Fourth, fertilization of the oocyte by X-bearing sperm versus Y-bearing sperm, and therefore the sex of the resulting zygote, is assigned with equal probabilities. This procedure assumes a chromosomal sex determination system with mendelian segregation of X and Y chromosomes in males.

Fifth, zygotes are randomly classified as viable or inviable based on sex-specific, marker-associated viability effects estimated from the extended $Cx(Co)^m$ model. This requires each zygote be randomly assigned one of two equally likely reciprocal marker complements, and the probability of viable/inviable classification is dependent on the marker complement assignment.

Finally, among the zygotes classified as viable conditional on reciprocal marker complement assignment, a second marker-independent round of viability selection was simulated. The probability of being reclassified as inviable was based on the empirical zygotic lethality rates estimated in marker-free control crosses in the accompanying paper.

This algorithm generates one data point classifiable as either one of the sex-specific phenotypic classes scored for recombination or an inviable zygote. Iteration of the process $n$ times to match number of eggs laid in a given dataset can be solved for the maximum likelihood solution of parameter values in the extended $Cx(Co)^m$ model, giving a single simulated estimate of the parameters from the data-generating process. Repeating this process, say 10,000 times, builds an empirical sampling distribution tailored to the experimental design of the dataset and allows construction of null hypothesis tests and confidence intervals.

#### Data availability

Further documentation of model structure and fitting procedures applied to the accompanying paper can be found at https://github.com/StevisonLab/CxCoM-with-Viability.

#### Acknowledgments

We would like to thank members of the Stevison Lab for discussion during the development of this project.

#### Funding

This research was supported by the National Institute of General Medical Sciences of the National Institutes of Health under award number R35GM147501. The content is solely the responsibility of the authors and does not necessarily represent the official view of the National Institutes of Health (NIH).

#### Conflicts of interest

The authors declare no conflicts of interest in reporting this research.
