## Supplemental Material for "An extension of the *Cx(Co)*^*m*^ model of crossover patterning to account for experimental mortality in *Drosophila melanogaster*"

**File S1. Supplementary material to "An extension of the *Cx(Co)$^m$* of crossover patterning to account for experimental mortality in *Drosophila melanogaster*."**

**Table of Contents:**


***I. Egg-to-adult viability*** ... **2**
*Figure S1. Standard curves for adjusting digital egg undercounts* ... *2*
*Table S1. Regression analysis of marker-free viability.* ... *3*
*Table S2. Regression analysis of multiply-marked viability.* ... *3*

***II. Single-locus analysis*** ... **4**
*Table S3. Single-locus analysis of the viability effects associated with scute.* ... *5*
*Table S4. Single-locus analysis of the viability effects associated with crossveinless.* ... *5*
*Table S5. Single-locus analysis of the viability effects associated with vermilion.* ... *6*
*Table S6. Single-locus analysis of the viability effects associated with forked.* ... *6*
*Table S7. Single-locus analysis of the viability effects associated with carnation* ... *6*
*Table S8. Single-locus analysis of the viability effects associated with intrachromosomal duplication Dp(1;1)sc$^{V1}$, y+.* ... *7*
*Figure S2. Single-locus, independently-estimated, sex-specific viability effects.* ... *7*

***III. Multi-locus analysis*** ... **8**
*Table S9. Multi-locus analysis of viability effects associated marker loci* ... *9*
*Figure S3. Multi-locus, simultaneously-estimated, sex-specific viability effects.* ... *10*

***IV. Alternate pooling scheme*** ... **11**
*Table S10. X chr. map length ANOVA of H3: Cx(Co)$^m$ pooling by individual vial.* ... *11*
*Table S11. X chr. map length ANOVA of H3: Cx(Co)$^m$ pooling by replicate cross.* ... *11*
*Table S12. X chr. map length ANOVA of H3: Cx(Co)$^m$ pooling by brooding period* ... *11*

***V. Stock Construction*** ... **12**
*A. Reconstitution of multiply-marked X chromosome strain* ... *12*
*B. Multiply-marked X chr. genetic background replacement* ... *13*
*C. Reconstitution of multiply-marked X chr. tester strain* ... *14*
*D. Multiply-marked X chr. genetic background replacement of tester strain* ... *15*
*E. Introgression of recessive X chr. marker into isogenic line* ... *16*
*F. Replacement of recessive X chr. marker in an isogenic line* ... *17*

**I. Egg-to-adult viability**

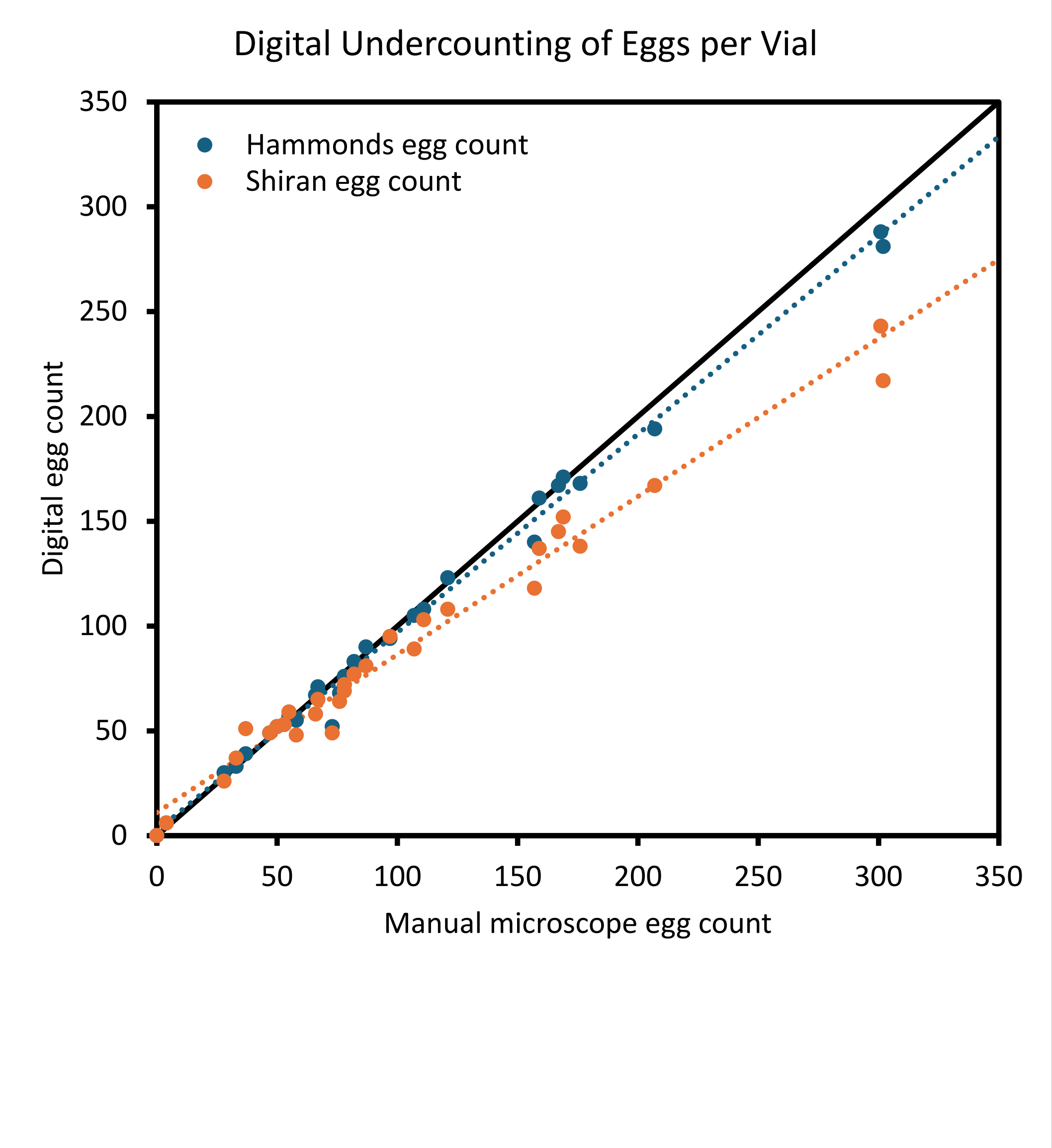


**Figure S1. Standard curves for adjusting digital egg undercounts.** Two independent digital egg counts were performed. Because digital scoring consistently undercounts eggs per vial and individual observers differ in their undercounting, both egg counts were transformed using standard curves constructed from independent dataset of Schneider *et al.* (2025). Illustrated is type II (major axis) regression of digital egg counts on manual microscope counts adjusted each line (blue dotted y = 0.9460x + 2.38 and orange dotted y =0.7537x + 11.07). Transformation of digital egg counts brings both dotted lines to match the 1:1 black solid line thereby adjusting for each observer's bias in digital egg detection.

Schneider, C., Koury, S., & Stevison, L. S. (2025). Protocol: An absolute egg-to-adult viability assay in *Drosophila melanogaster*. *Micropublication Biology*, *2025*, 10-17912.

**Table S1. Regression analysis of marker-free viability.** ANOVA table for comparison of male versus female egg-to-adult viability (assuming 1:1 primary sex ratio) in marker-free X chromosome control crosses.

| Source | df | SS | MS | F value | Pr(>F) |
|---|---|---|---|---|---|
| Brood | 1 | 0.05981488 | 0.05981488 | 2.39407245 | 0.12347767 |
| Sex | 1 | 0.01397553 | 0.01397553 | 0.55936634 | 0.45545019 |
| Cross | 9 | 0.29241155 | 0.03249017 | 1.30040941 | 0.23907265 |
| Residuals | 188 | 4.69709955 | 0.02498457 | | |

**Table S2. Regression analysis of multiply-marked viability.** ANOVA table for comparison of male versus female egg-to-adult viability (assuming 1:1 primary sex ratio) in multiply-marked X chromosome testcrosses.

| Source | df | SS | MS | F value | Pr(>F) |
|---|---|---|---|---|---|
| Brood | 1 | 0.01202625 | 0.01202625 | 1.53208958 | 0.218 |
| Sex | 1 | 0.56130645 | 0.56130645 | 71.5078622 | 5.35e-14 |
| Cross | 9 | 0.21221209 | 0.02357912 | 3.00387165 | 0.00278 |
| Residuals | 128 | 1.00474582 | 0.00784958 | | |

#### II. Single-locus analysis

The F2 experimental mortality in multiply-marked crosses is associated with visible markers used to score recombinant chromosomes. To model the uneven recovery of markers in both sexes we use four different 5-equation systems (*H0*, *H1*, *H2*, and *H3*). All models assume mendelian segregation of alleles in F1 females, and mendelian segregation of X and Y chromosomes in F1 males. The number of eggs laid is denoted $n$, and $v_x$ respresent the proportional viability due to random sources of experimental mortality. In order of increasing complexity these models are:

*H0*: no excess experimental mortality in multiply-marked crosses above and beyond the marker-free cross,

$$\begin{aligned}\text{female wildtype count} &= \tfrac{1}{4}nv_x\\ \text{male wildtype count} &= \tfrac{1}{4}nv_x\\ \text{female mutant count} &= \tfrac{1}{4}nv_x\\ \text{male mutant count} &= \tfrac{1}{4}nv_x\\ \text{inviable zygote count} &= n(1-v_x)\end{aligned}$$

*H1*: excess experimental mortality $v_1$ in multiply-marked crosses is random with respect to marker alleles,

$$\begin{aligned}\text{female wildtype count} &= \tfrac{1}{4}nv_xv_1\\ \text{male wildtype count} &= \tfrac{1}{4}nv_xv_1\\ \text{female mutant count} &= \tfrac{1}{4}nv_xv_1\\ \text{male mutant count} &= \tfrac{1}{4}nv_xv_1\\ \text{inviable zygote count} &= n(1-v_xv_1)\end{aligned}$$

*H2*: excess experimental mortality in multiply-marked crosses is solely due to mutant alleles at marker loci ($v_2$ for females and $v_3$ for males),

$$\begin{aligned}\text{female wildtype count} &= \tfrac{1}{4}nv_x\\ \text{male wildtype count} &= \tfrac{1}{4}nv_x\\ \text{female mutant count} &= \tfrac{1}{4}nv_xv_2\\ \text{male mutant count} &= \tfrac{1}{4}nv_xv_3\\ \text{inviable zygote count} &= \tfrac{1}{4}n\big(4-v_x(2+v_2+v_3)\big)\end{aligned}$$

*H3*: excess experimental mortality in multiply-marked crosses is due to viability effects associated with both mutant and wildtype alleles at marker loci ($v_4$, $v_5$, $v_6$, and $v_7$),

$$\text{female wildtype count} = \frac{1}{4} n v_x v_4$$
$$\text{male wildtype count} = \frac{1}{4} n v_x v_5$$
$$\text{female mutant count} = \frac{1}{4} n v_x v_6$$
$$\text{male mutant count} = \frac{1}{4} n v_x v_7$$
$$\text{inviable zygote count} = \frac{1}{4} n\big(4 - v_x(v_4 + v_5 + v_6 + v_7)\big)$$

The proportional viability due to random sources of experimental mortality $v_x = 0.921$.
All estimates of viability parameters are bounded by the inequalities
$H_1: 1 > v_1 > 0$,
$H_2: 1 > v_3 \neq v_4 > 0$, and
$H_3: 1 > v_1 \neq v_2 \neq v_3 \neq v_4 > 0$.

**Table S3. Single-locus analysis of the viability effects associated with *scute*.** All females are either heterozygous (Female +) or homozygous (Female -) while all males are hemizygous for wildtype (Male +) or mutant allele (Male -). *G*-tests with corresponding degrees of freedom and p-values provided for all viability models except *H3* which has 0 degrees of freedom to perform the test.

| | Female + | Male + | Female - | Male - | G_statistic | df | p_value |
|---|---|---|---|---|---|---|---|
| *H0:* | 1.000 | 1.000 | 1.000 | 1.000 | 10599.2821 | 4 | 0 |
| *H1:* | 0.610 | 0.610 | 0.610 | 0.610 | 256.407282 | 3 | 2.691e-55 |
| *H2:* | 1.000 | 1.000 | 0.398 | 0.236 | 1063.05147 | 2 | 1.450e-231 |
| *H3:* | 0.766 | 0.657 | 0.591 | 0.428 | NA | 0 | NA |

**Table S4. Single-locus analysis of the viability effects associated with *crossveinless*.** All females are either heterozygous (Female +) or homozygous (Female -) while all males are hemizygous for wildtype (Male +) or mutant allele (Male -). *G*-tests with corresponding degrees of freedom and p-values provided for all viability models except *H3* which has 0 degrees of freedom to perform the test.

| | Female + | Male + | Female - | Male - | G_statistic | df | p_value |
|---|---|---|---|---|---|---|---|
| *H0:* | 1.000 | 1.000 | 1.000 | 1.000 | 10965.321 | 4 | 0 |
| *H1:* | 0.610 | 0.610 | 0.610 | 0.610 | 622.447 | 3 | 1.371e-134 |
| *H2:* | 1.000 | 1.000 | 0.395 | 0.192 | 640.885 | 2 | 6.818e-140 |
| *H3:* | 0.817 | 0.748 | 0.540 | 0.337 | NA | 0 | NA |

**Table S5. Single-locus analysis of the viability effects associated with *vermilion*.** All females are either heterozygous (Female +) or homozygous (Female -) while all males are hemizygous for wildtype (Male +) or mutant allele (Male -). *G*-tests with corresponding degrees of freedom and p-values provided for all viability models except $H_3$ which has 0 degrees of freedom to perform the test.

| | Female + | Male + | Female - | Male - | G_statistic | df | p_value |
|---|---|---|---|---|---|---|---|
| *H0:* | 1.000 | 1.000 | 1.000 | 1.000 | 10646.201 | 4 | 0 |
| *H1:* | 0.610 | 0.610 | 0.610 | 0.610 | 303.326 | 3 | 1.896e-65 |
| *H2:* | 1.000 | 1.000 | 0.399 | 0.227 | 987.515 | 2 | 3.664e-215 |
| *H3:* | 0.772 | 0.673 | 0.584 | 0.412 | NA | 0 | NA |

**Table S6. Single-locus analysis of the viability effects associated with *forked*.** All females are either heterozygous (Female +) or homozygous (Female -) while all males are hemizygous for wildtype (Male +) or mutant allele (Male -). *G*-tests with corresponding degrees of freedom and p-values provided for all viability models except $H_3$ which has 0 degrees of freedom to perform the test.

| | Female + | Male + | Female - | Male - | G_statistic | df | p_value |
|---|---|---|---|---|---|---|---|
| *H0:* | 1.000 | 1.000 | 1.000 | 1.000 | 10501.470 | 4 | 0 |
| *H1:* | 0.610 | 0.610 | 0.610 | 0.610 | 158.595 | 3 | 3.684e-34 |
| *H2:* | 1.000 | 1.000 | 0.404 | 0.252 | 1289.474 | 2 | 9.868e-281 |
| *H3:* | 0.739 | 0.619 | 0.617 | 0.466 | NA | 0 | NA |

**Table S7. Single-locus analysis of the viability effects associated with *carnation*.** All females are either heterozygous (Female +) or homozygous (Female -) while all males are hemizygous for wildtype (Male +) or mutant allele (Male -). *G*-tests with corresponding degrees of freedom and p-values provided for all viability models except $H_3$ which has 0 degrees of freedom to perform the test.

| | Female + | Male + | Female - | Male - | G_statistic | df | p_value |
|---|---|---|---|---|---|---|---|
| *H0:* | 1.000 | 1.000 | 1.000 | 1.000 | 10486.643 | 4 | 0 |
| *H1:* | 0.610 | 0.610 | 0.610 | 0.610 | 143.769 | 3 | 5.819e-31 |
| *H2:* | 1.000 | 1.000 | 0.398 | 0.261 | 1331.266 | 2 | 8.302e-290 |
| *H3:* | 0.741 | 0.606 | 0.615 | 0.479 | NA | 0 | NA |

**Table S8. Single-locus analysis of viability effects associated with intrachromosomal duplication *Dp(1;1)sc$^{V1}$, y+*.** All females are either heterozygous (Female +) or homozygous (Female -) while all males are hemizygous for wildtype (Male +) or mutant allele (Male -). *G*-tests with corresponding degrees of freedom and p-values provided for all viability models except $H_3$ which has 0 degrees of freedom to perform the test.

| | Female + | Male + | Female - | Male - | G_statistic | df | p_value |
|---|---|---|---|---|---|---|---|
| H0: | 1.000 | 1.000 | 1.000 | 1.000 | 10483.745 | 4 | 0 |
| H1: | 0.610 | 0.610 | 0.610 | 0.610 | 140.870 | 3 | 2.454e-30 |
| H2: | 1.000 | 1.000 | 0.400 | 0.260 | 1342.392 | 2 | 3.186e-292 |
| H3: | 0.738 | 0.606 | 0.619 | 0.479 | NA | 0 | NA |

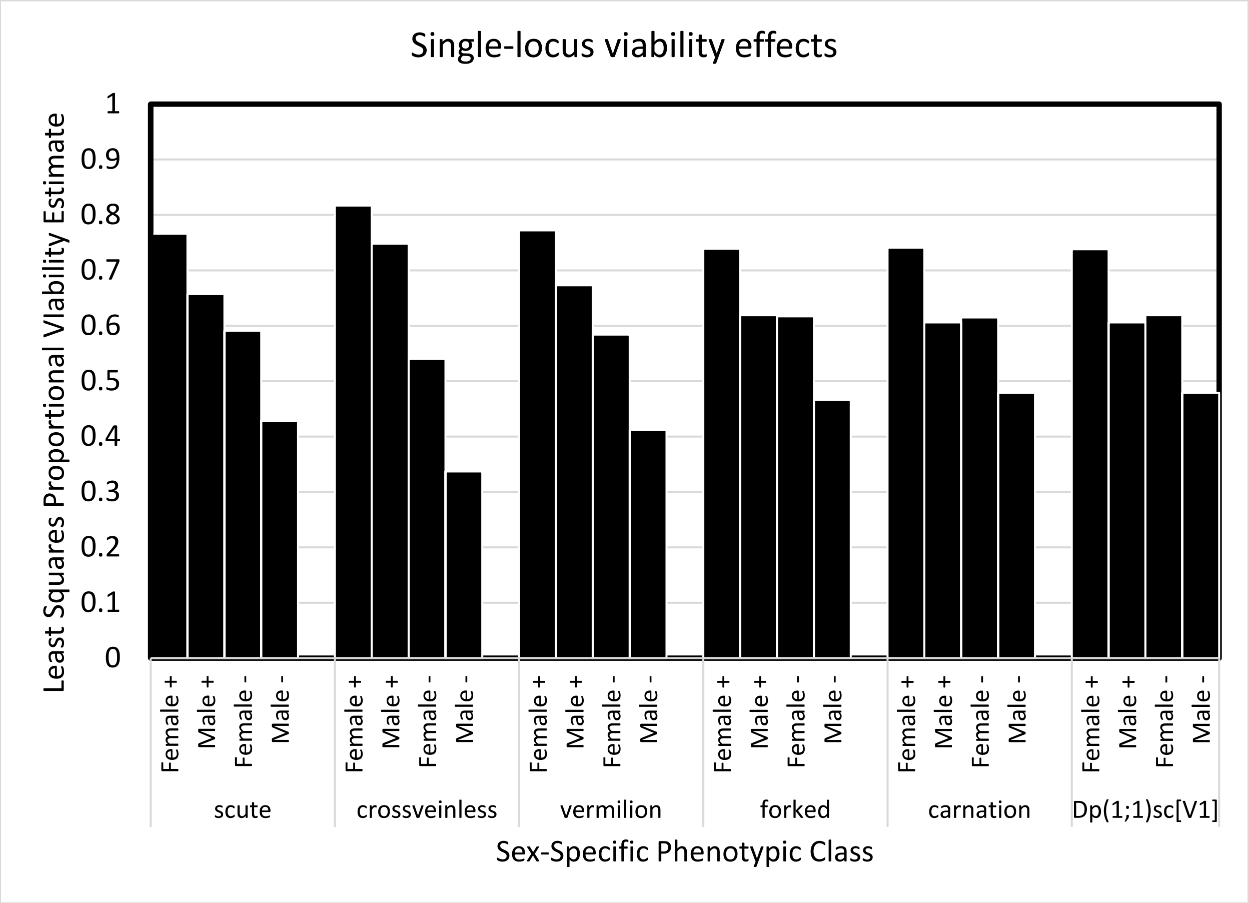


**Figure S2. Single-locus, independently-estimated, sex-specific viability effects.** Loci are listed in conventional order of *Drosophila melanogaster* X chromosome (left arm telomere to centromere). All females are either heterozygous (Female +) or homozygous (Female -) while all males are hemizygous for wildtype (Male +) or mutant allele (Male -). Please see main text for description of confounding effects of performing viability analysis independently on genetically linked loci.

#### III. Multi-locus analysis

As stated earlier in the single-locus analysis section, the $F_2$ experimental mortality in multiply-marked crosses is associated with visible markers used to score recombinant chromosomes. Because the six marker loci do not segregate independently, the single-locus tests confound each other and distort viability estimates by a factor related to the unknown genetic distances to be estimated. This problem can be solved by simultaneously estimating multi-locus marker viability effects, genetic lengths, and crossover interference by maximizing the likelihood of observed $F_2$ phenotypic class counts under the extended $Cx(Co)^m$ counting model. For a detailed description of this model please see Appendix. Here, we outline the constraints on viability estimates denoted $v_{alh}$ where $a$ is the inherited allele from the $F_1$ female (0 for wildtype and 1 for mutant), $l$ is the locus in a 6-point testcross, and $h$ is the sex of the $F_2$ progeny ($f$ for female and $m$ for male). These constraints are given in relation to the hypotheses in increasing complexity:

*$H_0$*: no excess experimental mortality in multiply-marked crosses above and beyond the marker-free cross,

$$1 = v_{01f} = v_{01m} = v_{11f} = v_{11m}$$
$$1 = v_{02f} = v_{02m} = v_{12f} = v_{12m}$$
$$1 = v_{03f} = v_{03m} = v_{13f} = v_{13m}$$
$$1 = v_{04f} = v_{04m} = v_{14f} = v_{14m}$$
$$1 = v_{05f} = v_{05m} = v_{15f} = v_{15m}$$
$$1 = v_{06f} = v_{06m} = v_{16f} = v_{16m}$$

*$H_1$*: excess experimental mortality in multiply-marked crosses is random with respect to marker alleles,

$$1 \geq v_{01f} = v_{01m} = v_{11f} = v_{11m}$$
$$1 \geq v_{02f} = v_{02m} = v_{12f} = v_{12m}$$
$$1 \geq v_{03f} = v_{03m} = v_{13f} = v_{13m}$$
$$1 \geq v_{04f} = v_{04m} = v_{14f} = v_{14m}$$
$$1 \geq v_{05f} = v_{05m} = v_{15f} = v_{15m}$$
$$1 \geq v_{06f} = v_{06m} = v_{16f} = v_{16m}$$

*$H_2$*: excess experimental mortality in multiply-marked crosses is solely due to mutant alleles at marker loci,

$$1 = v_{01f} = v_{01m} \geq v_{11f} \neq v_{11m}$$
$$1 = v_{02f} = v_{02m} \geq v_{12f} \neq v_{12m}$$
$$1 = v_{03f} = v_{03m} \geq v_{13f} \neq v_{13m}$$
$$1 = v_{04f} = v_{04m} \geq v_{14f} \neq v_{14m}$$
$$1 = v_{05f} = v_{05m} \geq v_{15f} \neq v_{15m}$$
$$1 = v_{06f} = v_{06m} \geq v_{16f} \neq v_{16m}$$

*H3*: excess experimental mortality in multiply-marked crosses is due to viability effects associated with both mutant and wildtype alleles at marker loci,

$$1 \geq v_{01f} \neq v_{01m} \neq v_{11f} \neq v_{11m}$$
$$1 \geq v_{02f} \neq v_{02m} \neq v_{12f} \neq v_{12m}$$
$$1 \geq v_{03f} \neq v_{03m} \neq v_{13f} \neq v_{13m}$$
$$1 \geq v_{04f} \neq v_{04m} \neq v_{14f} \neq v_{14m}$$
$$1 \geq v_{05f} \neq v_{05m} \neq v_{15f} \neq v_{15m}$$
$$1 \geq v_{06f} \neq v_{06m} \neq v_{16f} \neq v_{16m}$$

**Table S9. Multi-locus analysis of viability effects associated marker loci.** All females are either heterozygous or mutant homozygotes, while all males are hemizygous for wildtype or mutant allele. All values estimated simultaneously with genetic lengths and crossover interference using maximum likelihood methods with the extended *Cx(Co)^m* counting model.

| | Female Heterozygotes | | | | | |
|---|---|---|---|---|---|---|
| Hypothesis | *sc* | *cv* | *v* | *f* | *car* | *Dp(1:1)sc[V1]* |
| *H0:* | 1.00* | 1.00* | 1.00* | 1.00* | 1.00* | 1.00* |
| *H1:* | 1.00* | 1.00* | 1.00* | 1.00* | 1.00* | 1.00* |
| *H2:* | 1.00* | 1.00* | 1.00* | 1.00* | 1.00* | 1.00* |
| *H3:* | 0.99 | 1.00 | 0.88 | 1.00 | 1.00 | 1.00 |

| | Male Hemizygous Wildtype | | | | | |
|---|---|---|---|---|---|---|
| Hypothesis | *sc* | *cv* | *v* | *f* | *car* | *Dp(1:1)sc[V1]* |
| *H0:* | 1.00* | 1.00* | 1.00* | 1.00* | 1.00* | 1.00* |
| *H1:* | 1.00* | 1.00* | 1.00* | 1.00* | 1.00* | 1.00* |
| *H2:* | 1.00* | 1.00* | 1.00* | 1.00* | 1.00* | 1.00* |
| *H3:* | 0.88 | 1.00 | 1.00 | 1.00 | 0.89 | 1.00 |

| | Mutant Homozygotes | | | | | |
|---|---|---|---|---|---|---|
| Hypothesis | *sc* | *cv* | *v* | *f* | *car* | *Dp(1:1)sc[V1]* |
| *H0:* | 1.00* | 1.00* | 1.00* | 1.00* | 1.00* | 1.00* |
| *H1:* | 1.00* | 1.00* | 1.00* | 1.00* | 1.00* | 1.00* |
| *H2:* | 0.91 | 0.67 | 0.93 | 1.00 | 0.92 | 0.84 |
| *H3:* | 1.00 | 0.67 | 0.87 | 1.00 | 1.00 | 0.88 |

| | Hemizygous Mutant | | | | | |
|---|---|---|---|---|---|---|
| Hypothesis | *sc* | *cv* | *v* | *f* | *car* | *Dp(1:1)sc[V1]* |
| *H0:* | 1.00* | 1.00* | 1.00* | 1.00* | 1.00* | 1.00* |
| *H1:* | 1.00* | 1.00* | 1.00* | 1.00* | 1.00* | 1.00* |
| *H2:* | 0.94 | 0.43 | 0.93 | 0.89 | 1.00 | 0.77 |
| *H3:* | 1.00 | 0.43 | 1.00 | 0.90 | 1.00 | 0.83 |

* Proportional viability of 1.00 by assumption

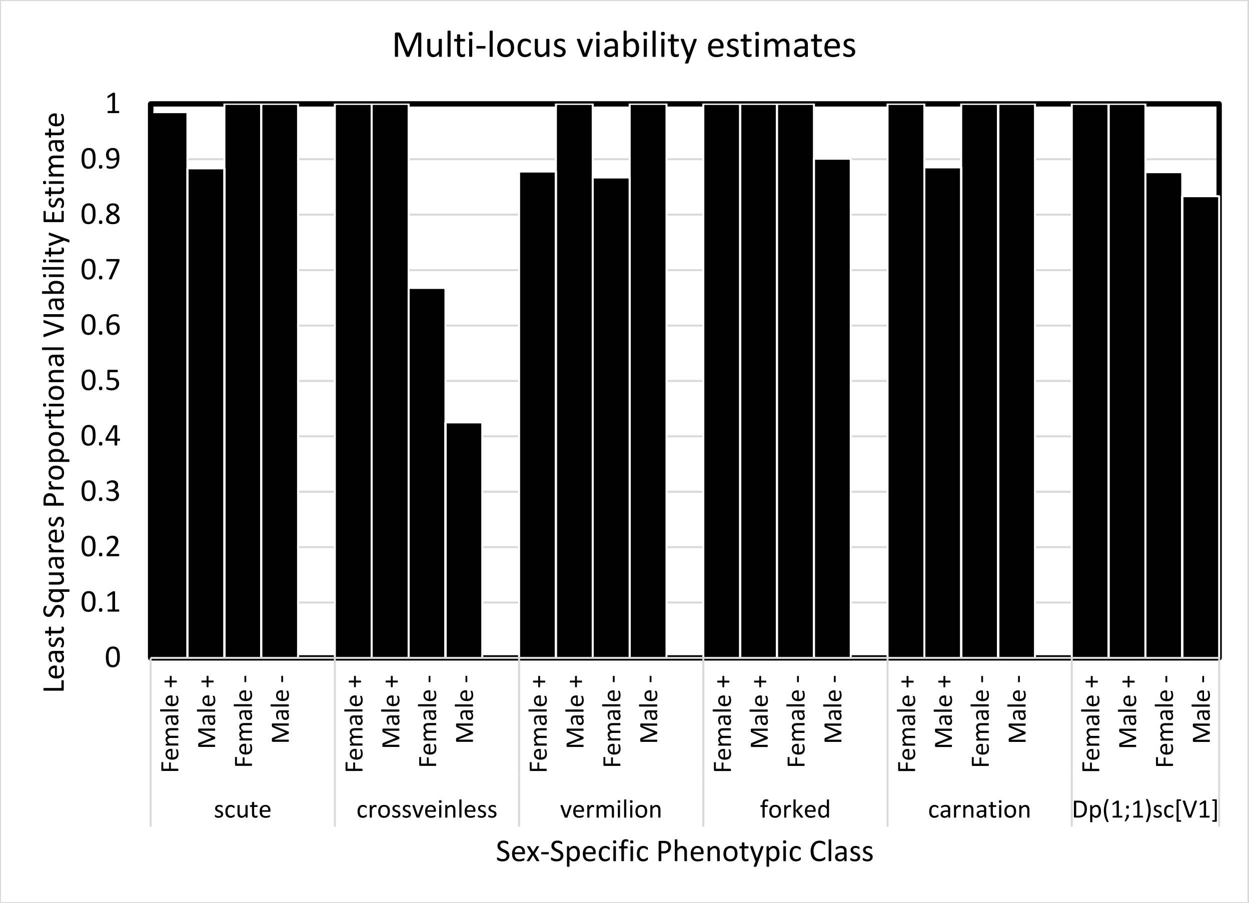


**Figure S3. Multi-locus, simultaneously-estimated, sex-specific viability effects.** Loci are listed in conventional order of *Drosophila melanogaster* X chromosome (left arm telomere to centromere). All females are either heterozygous (Female +) or homozygous (Female -) while all males are hemizygous for wildtype (Male +) or mutant allele (Male -). All estimates are from the full viability model $H_3$. Comparison with Figure S2 (based on single-locus $H_3$) demonstrates the strong confounding effects from *crossveinless* on viability estimates and reveals the negative viability effects linked to several wildtype alleles.

#### IV. Alternate pooling scheme

The same recombination dataset can be analyzed as a single experimental unit (as in prior sections of the supplemental material) or subdividing that dataset by the individual vials, replicate crosses, and/or brooding periods the $F_2$ data came from. We performed ANOVAs under these alternate pooling schemes to explore the variability in genetic length estimates. This procedure generated mean square error estimates that can be used for experimental design and constructing power curves (see main text figure 3).

**Table S10. X chromosome map length ANOVA of H3: *Cx(Co)$^m$* pooling by individual vial.** ANOVA table for total genetic length of the *Drosophila melanogaster* X chromosome when $F_2$ progeny are pooled by individual vials accounting for non-significant brood and cross effects.

| Source | Df | Sum Sq | Mean Sq | F value | Pr(>F) |
|---|---|---|---|---|---|
| Brood | 6 | 389.11073 | 64.8517883 | 1.09782989 | 0.37570133 |
| Cross | 9 | 554.846221 | 61.6495801 | 1.04362198 | 0.41862257 |
| Residuals | 54 | 3189.9264 | 59.0727112 | NA | NA |

**Table S11. X chromosome map length ANOVA of H3: *Cx(Co)$^m$* pooling by replicate cross.** ANOVA table for total genetic length of the *Drosophila melanogaster* X chromosome when $F_2$ progeny are pooled by replicated cross (fit as an intercept only model).

| Source | Df | Sum Sq | Mean Sq | F value | Pr(>F) |
|---|---|---|---|---|---|
| Residuals | 9 | 103.386067 | 11.4873408 | NA | NA |

**Table S12. X chr. map length ANOVA of H3: *Cx(Co)$^m$* pooling by brooding period.** ANOVA table for total genetic length of the *Drosophila melanogaster* X chromosome when $F_2$ progeny are pooled by brooding period (fit as an intercept only model).

| Source | Df | Sum Sq | Mean Sq | F value | Pr(>F) |
|---|---|---|---|---|---|
| Residuals | 6 | 35.3878444 | 5.89797406 | NA | NA |

#### V. Stock Construction

##### A. Reconstitution of multiply-marked X chromosome strain

### Starting Resources:

| | | |
|---|---|---|
| Marked X on uncontrolled background | $\frac{y+\ cv\ v\ f\ car}{y+\ cv\ v\ f\ car};\frac{+}{+};\frac{+}{+}$ | BDSC 1515 |
| Marked X on uncontrolled background | $\frac{y+++\ car{\cdot}y^{+}}{Y};\frac{+}{+};\frac{+}{+}$ | BDSC 4914 |
| Marked X on uncontrolled background | $\frac{y\ sc+\ +++}{Y};\frac{+}{+};\frac{+}{+}$ | BDSC 176 |

### Stable End Products:

| | |
|---|---|
| Multiply marked X chromosome | $\frac{y\ sc\ cv\ v\ f\ car{\cdot}y^{+}}{y\ sc\ cv\ v\ f\ car{\cdot}y^{+}};\frac{+}{+};\frac{+}{+}$ |
| Multiply marked X chromosome | $\frac{y\ sc\ cv\ v\ f\ car}{y\ sc\ cv\ v\ f\ car};\frac{+}{+};\frac{+}{+}$ |

### Crossing Scheme:

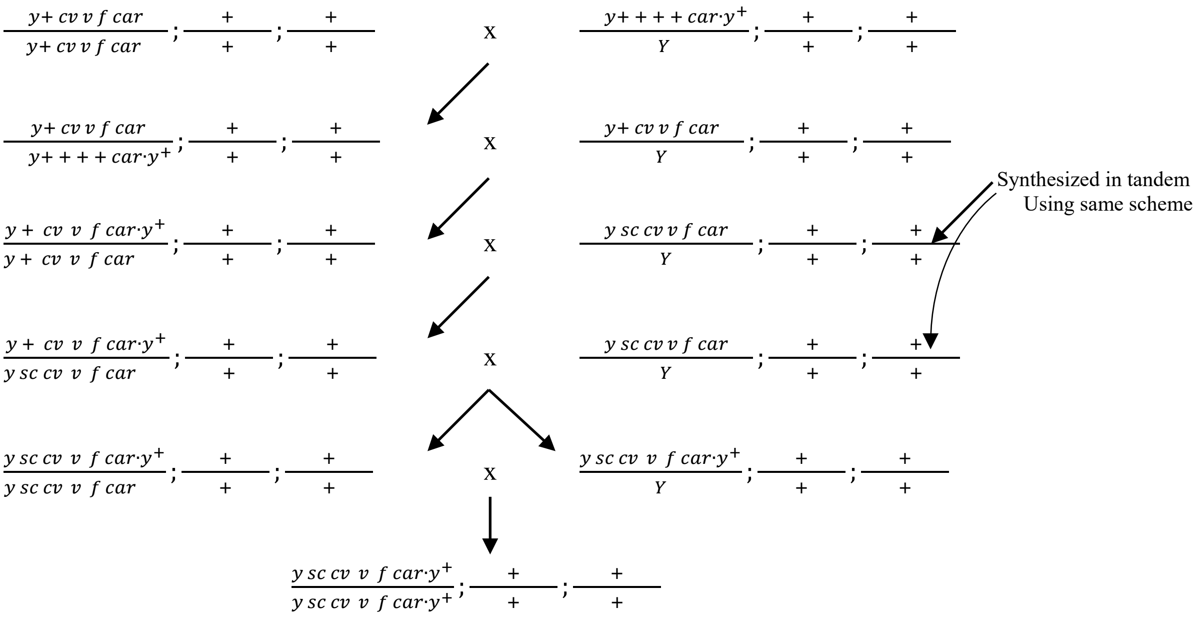

##### B. Multiply-marked X chr. genetic background replacement

### Starting Resources:

Marked X uncontrolled background $\frac{y\,sc\,cv\;\;v\;\;f\;car{\cdot}y^{+}}{y\,sc\,cv\;\;v\;\;f\;car{\cdot}y^{+}};\frac{+}{+};\frac{+}{+}$ See above

Isogenic source for genetic background $\frac{w^{1118}}{w^{1118}};\frac{BDSC\ 6326}{BDSC\ 6326};\frac{BDSC\ 6326}{BDSC\ 6326}$ BDSC 6326

Double balanced translocation stock $\frac{w^{*}}{Y};\frac{T(2;3)ap}{Ins(2L+2R)Cy;TM3}$ BDSC 2475

### Stable End Products:

Background replaced marked X chromosome $\frac{y\,sc\,cv\;\;v\;\;f\;car{\cdot}y^{+}}{y\,sc\,cv\;\;v\;\;f\;car{\cdot}y^{+}};\frac{BDSC\ 6326}{BDSC\ 6326};\frac{BDSC\ 6326}{BDSC\ 6326}$

Double balanced marked X chromosome $\frac{y\,sc\,cv\;\;v\;\;f\;car{\cdot}y^{+}}{y\,sc\,cv\;\;v\;\;f\;car{\cdot}y^{+}};\frac{T(2;3)ap}{Ins(2L+2R)Cy;TM3}$

### Crossing Scheme:

$\frac{y\,sc\,cv\;\;v\;\;f\;car{\cdot}y^{+}}{y\,sc\,cv\;\;v\;\;f\;car{\cdot}y^{+}};\frac{+}{+};\frac{+}{+}$ x $\frac{w^{*}}{Y};\frac{T(2;3)ap}{Ins(2L+2R)Cy;TM3}$

$\frac{y\,sc\,cv\;\;v\;\;f\;car{\cdot}y^{+}}{y\,sc\,cv\;\;v\;\;f\;car{\cdot}y^{+}};\frac{+}{+};\frac{+}{+}$ x $\frac{y\,sc\,cv\;\;v\;\;f\;car{\cdot}y^{+}}{Y};\frac{+}{Ins(2L+2R)Cy};\frac{+}{TM3}$

$\frac{y\,sc\,cv\;\;v\;\;f\;car{\cdot}y^{+}}{y\,sc\,cv\;\;v\;\;f\;car{\cdot}y^{+}};\frac{+}{Ins(2L+2R)Cy};\frac{+}{TM3}$ x $\frac{y\,sc\,cv\;\;v\;\;f\;car{\cdot}y^{+}}{Y};\frac{T(2;3)ap}{BDSC\ 6326;\ BDSC\ 6326}$

Synthesized the same way
Then outcross to *BDSC 6326*

$\frac{y\,sc\,cv\;\;v\;\;f\;car{\cdot}y^{+}}{y\,sc\,cv\;\;v\;\;f\;car{\cdot}y^{+}};\frac{BDSC\ 6326}{Ins(2L+2R)Cy};\frac{BDSC\ 6326}{TM3}$ x $\frac{y\,sc\,cv\;\;v\;\;f\;car{\cdot}y^{+}}{Y};\frac{BDSC\ 6326}{Ins(2L+2R)Cy};\frac{BDSC\ 6326}{TM3}$

$\frac{y\,sc\,cv\;\;v\;\;f\;car{\cdot}y^{+}}{y\,sc\,cv\;\;v\;\;f\;car{\cdot}y^{+}};\frac{BDSC\ 6326}{BDSC\ 6326};\frac{BDSC\ 6326}{BDSC\ 6326}$ x $\frac{y\,sc\,cv\;\;v\;\;f\;car{\cdot}y^{+}}{Y};\frac{BDSC\ 6326}{BDSC\ 6326};\frac{BDSC\ 6326}{BDSC\ 6326}$

$\frac{y\,sc\,cv\;\;v\;\;f\;car{\cdot}y^{+}}{y\,sc\,cv\;\;v\;\;f\;car{\cdot}y^{+}};\frac{BDSC\ 6326}{BDSC\ 6326};\frac{BDSC\ 6326}{BDSC\ 6326}$

##### C. Reconstitution of multiply-marked X chr. tester strain

### Starting Resources:

Marked X on uncontrolled background $\frac{y+\ cv\ v\ f\ car}{y+\ cv\ v\ f\ car};\frac{+}{+};\frac{+}{+}$ BDSC 1515

Marked X on uncontrolled background $\frac{y\ sc+ + + +}{Y};\frac{+}{+};\frac{+}{+}$ BDSC 176

### Stable End Products:

Marked X chromosome for outcrossing $\frac{y\ sc\ cv\ v\ f\ car}{y\ sc\ cv\ v\ f\ car};\frac{+}{+};\frac{+}{+}$

### Crossing Scheme:

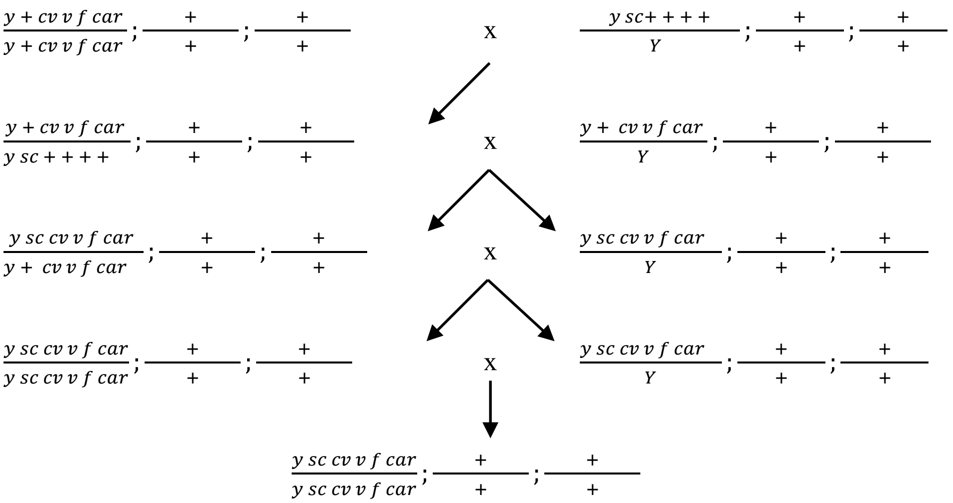

**D. Multiply-marked X chr. genetic background replacement of tester strain**

### Starting Resources:

Marked X uncontrolled background $\frac{y\ sc\ cv\ v\ f\ car}{y\ sc\ cv\ v\ f\ car};\frac{+}{+};\frac{+}{+}$

Isogenic source for genetic background $\frac{Canton-S}{Canton-S};\frac{Canton-S}{Canton-S};\frac{Canton-S}{Canton-S}$ BDSC 64349

Double balanced translocation stock $\frac{w^*}{Y};\frac{T(2;3)ap}{Ins(2L+2R)Cy;TM3}$ BDSC 2475

### Stable End Products:

Background replaced marked X chromosome $\frac{y\ sc\ cv\ v\ f\ car}{y\ sc\ cv\ v\ f\ car};\frac{Canton-S}{Canton-S};\frac{Canton-S}{Canton-S}$

Double balanced marked X chromosome $\frac{y\ sc\ cv\ v\ f\ car}{y\ sc\ cv\ v\ f\ car\cdot};\frac{T(2;3)ap}{Ins(2L+2R)Cy;TM3}$

### Crossing Scheme:

$\frac{y\ sc\ cv\ v\ f\ car}{y\ sc\ cv\ v\ f\ car};\frac{+}{+};\frac{+}{+}$ x $\frac{w^*}{Y};\frac{T(2;3)ap}{Ins(2L+2R)Cy;TM3}$

$\frac{y\ sc\ cv\ v\ f\ car}{y\ sc\ cv\ v\ f\ car};\frac{+}{+};\frac{+}{+}$ x $\frac{y\ sc\ cv\ v\ f\ car}{Y};\frac{+}{Ins(2L+2R)Cy};\frac{+}{TM3}$

$\frac{y\ sc\ cv\ v\ f\ car}{y\ sc\ cv\ v\ f\ car\cdot y^+};\frac{+}{Ins(2L+2R)Cy};\frac{+}{TM3}$ x $\frac{y\ sc\ cv\ v\ f\ car}{Y};\frac{T(2;3)ap}{Canton-S;\ Canton-S}$

Synthesized the same way
Then outcross to *Canton-S*

$\frac{y\ sc\ cv\ v\ f\ car}{y\ sc\ cv\ v\ f\ car};\frac{Canton-S}{Ins(2L+2R)Cy};\frac{Canton-S}{TM3}$ x $\frac{y\ sc\ cv\ v\ f\ car}{Y};\frac{Canton-S}{Ins(2L+2R)Cy};\frac{Canton-S}{TM3}$

$\frac{y\ sc\ cv\ v\ f\ car}{y\ sc\ cv\ v\ f\ car};\frac{Canton-S}{Canton-S};\frac{Canton-S}{Canton-S}$ x $\frac{y\ sc\ cv\ v\ f\ car}{Y};\frac{Canton-S}{Canton-S};\frac{Canton-S}{Canton-S}$

$\frac{y\ sc\ cv\ v\ f\ car}{y\ sc\ cv\ v\ f\ car};\frac{Canton-S}{Canton-S};\frac{Canton-S}{Canton-S}$

**E. Introgression of recessive X chr. marker into isogenic line**

### Starting Resources:

| | | |
|---|---|---|
| Isogenic source for genetic background | $\frac{DGRP\ 399}{DGRP\ 399};\frac{DGRP\ 399}{DGRP\ 399};\frac{DGRP\ 399}{DGRP\ 399}$ | BDSC 25192 |
| Marked X on uncontrolled background | $\frac{y\ cv\ v\ f\ car}{Y};\frac{+}{+};\frac{+}{+}$ | BDSC 1515 |

### Stable End Products:

Recessive X marker isogenic (>99%) stock $\frac{DGRP\ 399\ y}{DGRP\ 399\ y};\frac{DGRP\ 399}{DGRP\ 399};\frac{DGRP\ 399}{DGRP\ 399}$

### Crossing Scheme:

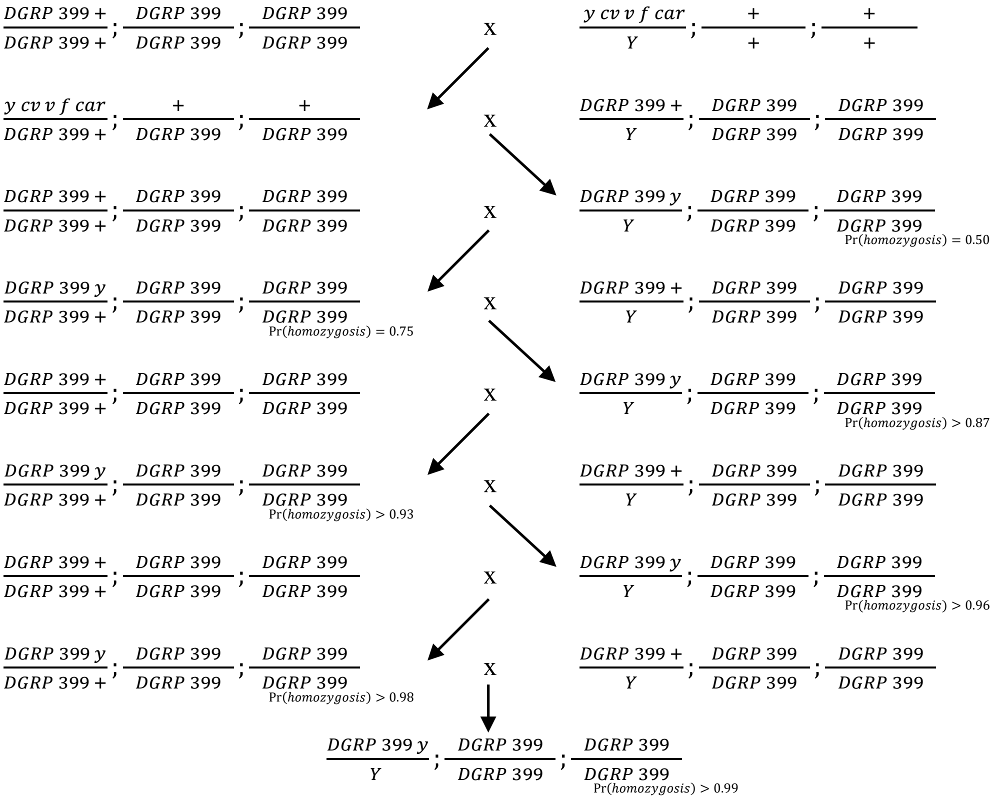

##### F. Replacement of recessive X chr. marker in an isogenic line

### Starting Resources:

| | | |
|---|---|---|
| Isogenic source for genetic background | $\frac{w^{1118}}{w^{1118}};\frac{BDSC\ 6326}{BDSC\ 6326};\frac{BDSC\ 6326}{BDSC\ 6326}$ | BDSC 6326 |
| Marked X on uncontrolled background | $\frac{w^{+}}{Y};\frac{Canton-S}{Canton-S};\frac{Canton-S}{Canton-S}$ | BDSC 64349 |

### Stable End Products:

| | |
|---|---|
| Recessive X marker isogenic (>99%) stock | $\frac{w^{+}}{w^{+}};\frac{BDSC\ 6326}{BDSC\ 6326};\frac{BDSC\ 6326}{BDSC\ 6326}$ |

### Crossing Scheme:

| | | |
|---|---|---|
| $\frac{w^{1118}}{w^{1118}};\frac{BDSC\ 6326}{BDSC\ 6326};\frac{BDSC\ 6326}{BDSC\ 6326}$ | X | $\frac{w^{+}}{Y};\frac{Canton-S}{Canton-S};\frac{Canton-S}{Canton-S}$ |
| $\frac{w^{1118}}{w^{+}};\frac{BDSC\ 6326}{Canton-S};\frac{BDSC\ 6326}{Canton-S}$ | X | $\frac{w^{1118}}{Y};\frac{BDSC\ 6326}{BDSC\ 6326};\frac{BDSC\ 6326}{BDSC\ 6326}$ |
| $\frac{w^{1118}}{w^{+}};\frac{BDSC\ 6326}{BDSC\ 6326};\frac{BDSC\ 6326}{BDSC\ 6326}$ $Pr(homozygosis) = 0.50$ | X | $\frac{w^{1118}}{Y};\frac{BDSC\ 6326}{BDSC\ 6326};\frac{BDSC\ 6326}{BDSC\ 6326}$ |
| $\frac{w^{1118}}{w^{+}};\frac{BDSC\ 6326}{BDSC\ 6326};\frac{BDSC\ 6326}{BDSC\ 6326}$ $Pr(homozygosis) = 0.75$ | X | $\frac{w^{1118}}{Y};\frac{BDSC\ 6326}{BDSC\ 6326};\frac{BDSC\ 6326}{BDSC\ 6326}$ |
| $\frac{w^{1118}}{w^{+}};\frac{BDSC\ 6326}{BDSC\ 6326};\frac{BDSC\ 6326}{BDSC\ 6326}$ $Pr(homozygosis) > 0.87$ | X | $\frac{w^{1118}}{Y};\frac{BDSC\ 6326}{BDSC\ 6326};\frac{BDSC\ 6326}{BDSC\ 6326}$ |
| $\frac{w^{1118}}{w^{+}};\frac{BDSC\ 6326}{BDSC\ 6326};\frac{BDSC\ 6326}{BDSC\ 6326}$ $Pr(homozygosis) > 0.93$ | X | $\frac{w^{1118}}{Y};\frac{BDSC\ 6326}{BDSC\ 6326};\frac{BDSC\ 6326}{BDSC\ 6326}$ |
| $\frac{w^{1118}}{w^{+}};\frac{BDSC\ 6326}{BDSC\ 6326};\frac{BDSC\ 6326}{BDSC\ 6326}$ $Pr(homozygosis) > 0.96$ | X | $\frac{w^{+}}{Y};\frac{BDSC\ 6326}{BDSC\ 6326};\frac{BDSC\ 6326}{BDSC\ 6326}$ $Pr(homozygosis) > 0.96$ |
| $\frac{w^{+}}{w^{+}};\frac{BDSC\ 6326}{BDSC\ 6326};\frac{BDSC\ 6326}{BDSC\ 6326}$ $Pr(homozygosis) > 0.98$ | X | $\frac{w^{+}}{Y};\frac{BDSC\ 6326}{BDSC\ 6326};\frac{BDSC\ 6326}{BDSC\ 6326}$ $Pr(homozygosis) > 0.98$ |

$$\frac{w^{+}}{w^{+}};\frac{BDSC\ 6326}{BDSC\ 6326};\frac{BDSC\ 6326}{BDSC\ 6326}$$
$Pr(homozygosis) > 0.99$
